# The MEK1/2 pathway as a therapeutic target in high-grade serous ovarian carcinoma

**DOI:** 10.1101/772061

**Authors:** Mikhail Chesnokov, Imran Khan, Yeonjung Park, Jessica Ezel, Geeta Mehta, Abdelrahman Yousif, Linda J. Hong, Ronald J. Buckanovich, Ilana Chefetz

**Author notes:** **Correspondence:** Ilana Chefetz, PhD, The Hormel Institute, University of Minnesota, 801 16^th^ Ave NE Austin MN 55912.

## Abstract

**Rationale:** High-grade serous ovarian carcinoma (HGSOC) is the deadliest of gynecological cancers due to high rate of recurrence and acquired chemoresistance. Mutation and activation of the RAS/MAPK pathway has been linked to cancer cell proliferation and therapeutic resistance in numerous cancers. While RAS mutations are not commonly observed in HGSOC, less is known about downstream pathway activation. We therefore sought to investigate the role of MEK1/2 signaling in ovarian cancer.

**Methods:** MEK1/2 pathway activity was evaluated in clinical HGSOC tissue samples and ovarian cancer cell lines by using tissue microarray-based immunohistochemistry, immunoblotting, and RT-qPCR. OVCAR8 and PEO4 HGSOC cell lines were used to assess the effect of MEK1/2 inhibition on cell viability, proliferation rate, and stem-like characteristics. Xenografts were used in mice to investigate the effect of MEK1/2 inhibition on tumor growth *in vivo*. A drug washout experimental model was used to study the lasting effects of MEK1/2 inhibition therapy.

**Results:** MEK1/2 signaling is active in a majority of HGSOC tissue samples and cell lines. MEK1/2 is further stimulated by cisplatin treatment, suggesting that MEK1/2 activation may play a role in chemotherapy resistance. The MEK1/2 inhibitor, trametinib, drastically inhibits MEK1/2 downstream signaling activity, causes prominent cell cycle arrest in the G_1/0_-phase in cell cultures, and reduces the rate of tumor growth *in vivo*, but does not induce cell death. Cells treated with trametinib display a high CD133^+^ fraction and increased expression of stemness-associated genes. Transient trametinib treatment causes long-term increases in a high ALDH1 activity subpopulation of cells that possess the capability of surviving and growing in non-adherent conditions.

**Conclusions:** MEK1/2 inhibition in HGSOC cells efficiently inhibits proliferation and tumor growth and therefore may be a promising approach to suppress ovarian cancer cell growth. MEK1/2 inhibition promotes stem-like properties, thus suggesting a possible mechanism of resistance and that a combination with CSC-targeting drugs should be considered.

## Introduction

In the US, ovarian cancer ranks 5^th^ in cancer-related deaths in women, displaying the 3^rd^ highest mortality rate [1, 2]. About 90% of ovarian cancer cases are epithelial in origin [3] with high grade serous ovarian cancer (HGSOC) being the most common and deadly subtype [2, 4–6]. While most HGSOC tumors initially respond well to platinum-based therapy, about 75% of patients experience disease relapse [4, 7–10] due to acquired resistance to chemotherapeutic agents [4, 5, 11, 12]. Multiple mechanisms of chemoresistance include activation of DNA repair systems, increased drug efflux due to *ABCB1* membrane transporter overexpression, changes in drug-specific metabolism, and apoptosis inhibition [11, 13, 14].

Tumor heterogeneity likely contributes to chemotherapy resistance. Ovarian tumors are heterogeneous on both genomic and cellular levels [15, 16]. At the cellular level, analysis of cisplatin-sensitive vs. cisplatin-resistant tumors revealed that resistant cells descend from pre-existing minor subpopulations in the primary tumor [17], which most likely represent cancer stem-like cells (CSCs) [18–24]. CSCs possess high resistance to cytotoxic and genotoxic effects, are capable of self-renewal, and asymmetric division that generates a progeny of fast-proliferating bulk tumor cells. CSCs possess a high tumorigenic potential and can re-initiate tumor development after chemotherapeutic treatment [19, 25, 26]. Survival and proliferation of CSCs in various tumors has been shown to be highly dependent on activity of mitogen-activated protein kinase (MAPK) signaling [27–30].

The MEK1/2 branch of the MAPK pathway mainly stimulates cell proliferation, migration, and differentiation (Fig. 1A), while p38- and JNK/SAPK-associated signaling induces apoptosis, inflammation, and stress responses [31]. MEK1/2 signaling hyperactivation frequently occurs in malignant tumors, and is therefore a promising target for anticancer therapy [32, 33]. Because MEK1/2 selectively activates ERK1/2, its inhibition is an efficient way to suppress the activity of the whole cascade [34]. Based on clinical studies [35], MEK1/2 inhibitors received FDA approval for tumors with activating *BRAF* mutations, including melanoma (trametinib, cobimetinib and binimetinib), non-small cell lung cancer (trametinib), and thyroid cancer (trametinib) [36].

**Fig. 1.**
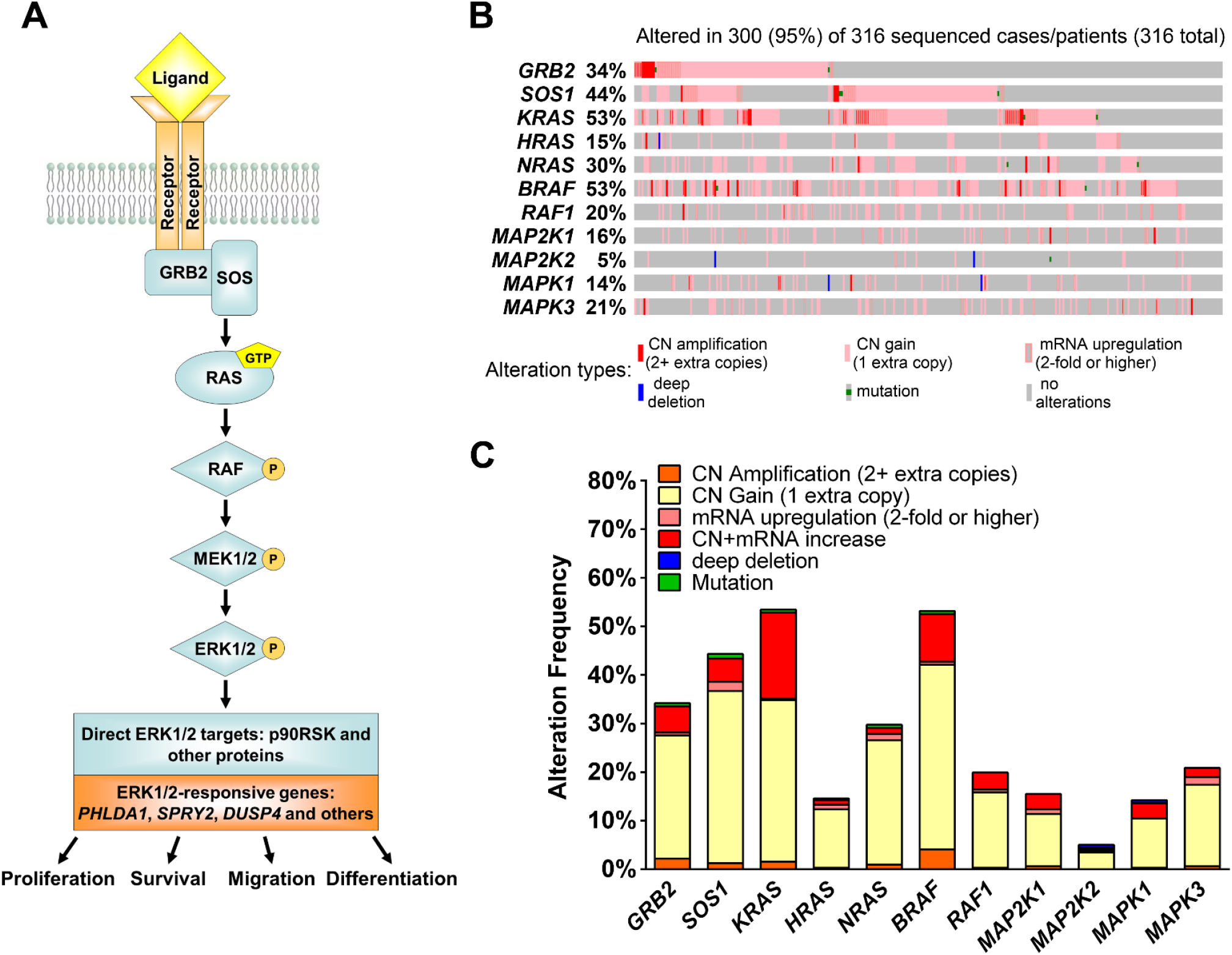
The MEK1/2 signaling pathway and its genetic alterations observed in HGSOC tissues. (A) The main elements of the MEK1/2 signaling pathway and cell properties controlled by its activity. (B) Genetic alterations and expression changes observed for MEK1/2 pathway elements in HGSOC tissues according to TCGA data. (C) Frequency of different genetic and transcriptional alterations of MEK1/2 pathway elements in HGSOC tissues. CN – copy number.

The function of the MEK1/2-ERK1/2 pathway has been mainly studied in low-grade ovarian tumors due to high frequency of activating *KRAS* and *BRAF* mutations [37, 38]. Despite the absence of *KRAS/BRAF* mutations in HGSOC, MEK1/2 signaling hyperactivation was reported in HGSOC and is associated with poor prognosis [39]. The present paper aims to investigate the role of high MEK1/2 activity in the regulation of proliferation, viability, and stemness of HGSOC cells.

## Materials and Methods

### Clinical tissue samples

Patient samples were obtained in accordance with protocols approved by the University of Michigan’s IRB (HUM0009149). Tumors were processed for protein isolation as previously described [23]. Tissue microarray (TMA) slides were constructed using paired clinical samples of HGSOC and normal fallopian tube tissue obtained from 10 patients by the Pathology Department core, Rhode Island Hospital, Brown University (LOCATION).

### Cell culture

OVCAR8, PEO4, and A2780 cells were provided by Dr. S. Murphy (Duke University). Kuramochi and OVSAHO cell lines were purchased from the Japanese cell line bank (LOCATION). TOV21D, HEY1, PEO1, and TOV21G cells were purchased from ATCC and Sigma. OVCAR8, PEO4, Kuramochi, OVSAHO, and PEO1 are HGSOC cell lines, whereas A2780, TOV21D, HEY1, and TOV21G cells are Type I ovarian cancer. All cell lines were cultivated in RPMI-1640 medium (Corning, USA) supplemented with 5% FBS (Gibco, USA) and penicillin/streptomycin (Corning, USA). PEO1 cells were cultivated in medium with the addition of 1 mM sodium pyruvate (Corning, USA).

### Drug treatment of cultured cells: (all sources need CITY and STATE)

Compounds used included trametinib (Selleck Chemicals, USA), Z-VAD-FMK (UBP Bio, USA), Nec-1 (ApexBio, USA), cisplatin (ApexBio, USA), and staurosporine (ApexBio, USA). Trametinib, Z-VAD-FMK, Nec-1, and staurosporine were dissolved in DMSO and cisplatin was dissolved in sterile water. Control samples in all experiments performed were treated with vehicle only. Vehicle concentration in growth medium did not exceed 0.2%.

### Western blotting

Total protein extracts were obtained from cell or tissue samples using Pierce RIPA ←define buffer (ThermoFisher Scientific, CITY, STATE, USA) according to manufacturer’s protocol. Protein concentrations were estimated using Pierce BCA Protein Assay Kit (ThermoFisher Scientific). Forty μg of total protein were separated in Bolt 4-12% Bis-Tris Plus Gels (ThermoFisher Scientific) and transferred to Hybond P 0.45 PVDF membranes (GE Healthcare, USA). Membranes were blocked with 5% BSA (Fisher Scientific, USA) in tris-buffered saline with Tween-20 (TBST) (Fisher Scientific, USA) and probed overnight at 4°C with the following primary antibodies diluted in 5% BSA in TBST: pMEK1/2 (CST 41G9, 1:1000), pERK1/2 (CST D13.14.4E, 1:1000), total ERK1/2 (CST 137F5, 1:1000), pp90RSK1 (R&D 1024A, 1:1000), GAPDH (ProteinTech 1E6D9, 1:10000), pSMAD2 (CST 138D4, 1:1000), pSTAT3 (CST D3A7, 1:1000), pGSK3b (CST D85E12, 1:1000), pCRAF (CST 56A6, 1:1000). Secondary antibodies were HRP-conjugated anti-rabbit IgG (CST 7074, 1:10000) or anti-mouse IgG (CST 7076, 1:10000) diluted in 5% skim milk (MilliporeSigma, USA) in TBST. Protein bands were developed using Luminata Classico or Luminata Forte HRP substrate (MilliporeSigma, USA) and detected using Amersham Imager 600 (GE Healthcare, USA).

### Immunohistochemical staining of TMA slides

TMA slides were deparaffinized and rehydrated according to common protocols. Antigen retrieval was performed by boiling the slides in citrate buffer (pH = 6.0) for 10 minutes using a microwave oven. Sections were blocked using solution of 10% FBS (Gibco, USA) and 1% BSA in TBS for 2 hours at room temperature and probed with pERK1/2 antibodies (CST D13.14.4E, 1:100 in TBS with 1% BSA) overnight at 4°C. Slides were rinsed with 0.0025% Triton-X100 (Fisher Scientific, USA) in TBS, probed with secondary HRP-conjugated anti-rabbit IgG (CST 7074, 1:1000 in TBS with 1% BSA) for 1 hour at room temperature and developed with a 3,3-diaminobenzidine kit (Vector Laboratories, USA). Separate slides were counterstained with hematoxylin (Fisher Scientific, USA). After staining slides were rinsed with DI water, dehydrated and mounted using toluene (Fisher Scientific, USA).

### RT-qPCR analysis of gene expression

Total RNA was isolated from cell or tissue samples using TRIzol reagent and the PureLink RNA Mini Kit (ThermoFisher Scientific, USA) with additional on-column DNAse treatment. Reverse transcription was performed using the RevertAid RT Reverse Transcription Kit (ThermoFisher Scientific, USA). Quantitative PCR analysis of gene expression levels was performed in a CFX96 Touch thermocycler (Bio-Rad Laboratories, USA) using PowerUp SYBR Green Master Mix (ThermoFisher Scientific, USA) and primers from Table S1. A three-step amplification program (15 seconds at 95°C, 45 seconds at 62°C, 30 seconds at 72°C) was run for 40 cycles and reaction specificity was checked by melt curve analysis and agarose electrophoresis. Reaction efficiency was evaluated using standard curve approach and was within 98-102% for all primers. Transcript abundance was estimated using Pfaffl’s method [40] *TBP* was used as a housekeeping normalization gene.

### Cell viability assay

Cells were plated in 12-well plates (Olympus Plastics, USA) at 100,000 cells/well. After 24 hours, growth medium was replaced with fresh medium containing compounds of interest. If Z-VAD-FMK or Nec-1 were used in treatment, cells were pre-treated with aforementioned compounds for 45 minutes before adding other compounds. After 72 hours of treatment, cells were harvested by trypsinization, pelleted, and resuspended in PBS. Numbers of viable and dead cells were assessed by direct counting using a Countess automated cell counter (ThermoFisher Scientific, USA) in the presence of 0.4% Trypan Blue. IC_50_ values were estimated based on relative viable cell numbers obtained for cells treated with different concentrations of cisplatin or trametinib.

### Spheroid cell growth assay

Cells were seeded in 24-well ultra-low attachment plates (Corning, USA) at 2,000 cells/well. After 7 days, cell clusters were harvested and disrupted by mild trypsinization, pelleted, and resuspended in PBS. Numbers of viable and dead cells were assessed by direct counting using a Countess automated cell counter (ThermoFisher Scientific, USA) in the presence of 0.4% Trypan Blue.

### Real-time cytotoxicity assay

Cells were plated in 96-well plates (Olympus Plastics, USA) at 5,000 cells/well. After 24 hours, growth medium was replaced with fresh medium containing compounds of interest and Cytotox Green Reagent (Bio-Essen, USA). Dead cells were detected in real time for 72 hours using the IncuCyte S3 cell imaging system (BioEssen, USA). The relative cytotoxicity level was estimated as the number of green fluorescent objects normalized to corresponding cell confluence values. Staurosporine (200 nM) was used as a positive control to induce cell death.

### Cell cycle assay

Cells were plated in 12-well plates at 100,000 cells/well. After 24 hours, growth medium was replaced with fresh medium containing compounds of interest. After 24 hours of treatment, cells were harvested by trypsinization, resuspended in 300 μL of ice-cold PBS, and fixed by the addition of 0.7 mL of ice-cold 70% ethanol in a dropwise manner with constant mixing. After addition of ethanol, samples were stored at −70°C overnight. Fixed cell samples were washed with ice-cold ethanol twice, treated with 0.2 mg/mL RNAse A for 60 minutes at 37°C, and stained with 10 μg/mL propidium iodide. Stained samples were analyzed using the FACSCalibur flow cytometer (BD Biosciences, USA) and ModFit LT software (Verity Software House, USA).

### Estimation of CD133^+^ cell fraction

Cells were harvested by trypsinization, washed with cold PBS, resuspended in PBS and stained with anti-CD133 antibodies conjugated with APC fluorophore (Miltenyi Biotec 293C3, 1:50) for 30 minutes at 4°C. Stained cells were washed with cold PBS and resuspended in PBS containing 0.1 μg/mL DAPI to exclude dead cells from analysis. Samples were analyzed using CytoFLEX flow cytometer and CytExpert software (Beckman Coulter, USA) according to previously described algorithms [23].

### Estimation of ALDH activity

Cells were harvested by trypsinization, washed with cold PBS and stained using the ALDEFLUOR Kit (STEMCELL Technologies, Canada) according to a previously described protocol [23]. Stained cells were resuspended in ALDEFLUOR Assay Buffer containing 0.1 ug/mL DAPI to exclude dead cells from analysis. Samples were analyzed using the CytoFLEX flow cytometer and CytExpert software (Beckman Coulter, USA). Cells displaying ALDEFLUOR signal at least 10-fold higher than median values were considered “high-positive” and their fraction was evaluated separately.

### Animal studies

All experiments were performed with approval of the University Committee on Use and Care of Animals at the University of Michigan. PEO-4 cells (100,000) in 100 μl of Matrigel (BD Biosciences, USA) were injected subcutaneously into the axillae of 8-week-old female NOD.Cg-Prkdcscid Il2rgtm1Wjl/SzJ (NSG) mice. Three days after cell injection, the mice were treated with IP injections of DMSO or 1 mg/kg trametinib daily (n = 10 mice per treatment group) for 25 days. Tumors were measured using calipers, and tumor volume (L x W x W/2) was calculated. After 25 days, mice were sacrificed, and tumor tissue samples were collected for analysis.

### Statistical analysis

At least 3 independent replicates were performed for each experiment. Differences between sample groups were evaluated using Mann-Whitney U-test or Kruskal-Wallis H-test. Statistical significance was accepted with *p* < 0.05. Asterisks, “*”, “**”, and “***” denote *p* < 0.05, *p* < 0.01, and *p* < 0.001, respectively. Half-maximal inhibitory concentration (IC_50_) and 50% growth inhibitory concentration (GI_50_) values were calculated using the nonlinear regression algorithm in Prism 7 software (GraphPad Software, USA). Comparison of IC_50_ and GI_50_ values was performed based on confidence intervals.

## Results

### The MEK1/2 pathway is active in high grade ovarian tumors

Activation of MEK1/2 signaling frequently occurs in cancer cells and promotes cell proliferation [32]. The MEK1/2 pathway has a clear hierarchical structure of signal transduction (Fig. 1A, Table S2). We used data available from The Cancer Genome Atlas database to analyze genetic and transcriptional changes in MEK1/2 pathway components occurring in HGSOC [41, 42]. While *KRAS* and *BRAF* mutations only occur in 1.26% of HGSOC cases, 25% of cases display amplifications (2 or more extra copies) in one or more of genes involved in MAPK signal transduction (Table S2). Taking local genetic gains (1 extra copy) and mRNA overexpression events into account increases the percentage of cases with pro-active changes in the MEK1/2 pathway to 95% (Fig. 1B). *SOS1, KRAS*, and *BRAF* are more often affected by these alterations compared to their downstream targets, MEK and ERK (Fig. 1C). Deletions of MEK-related genes are very rare in HGSOC comprising 1.6% of all cases (Fig. 1B).

To confirm activity at the protein level, we next analyzed the level of phosphorylated and total ERK1/2 (pERK1/2 and tERK1/2, respectively) in 43 clinically obtained HGSOC tumors. We detected pERK1/2 bands in 84% (36 of 43 cases) of samples by using a Western analysis (Fig. 2A). This observation was further confirmed using TMA slides constructed from paired samples of HGSOC and benign fallopian tube obtained from 10 independent patient samples. IHC analysis revealed intense pERK1/2 staining in all tumor samples similar or more prominent than in the corresponding normal samples (Fig. 2B, S1).

**Fig. 2.**
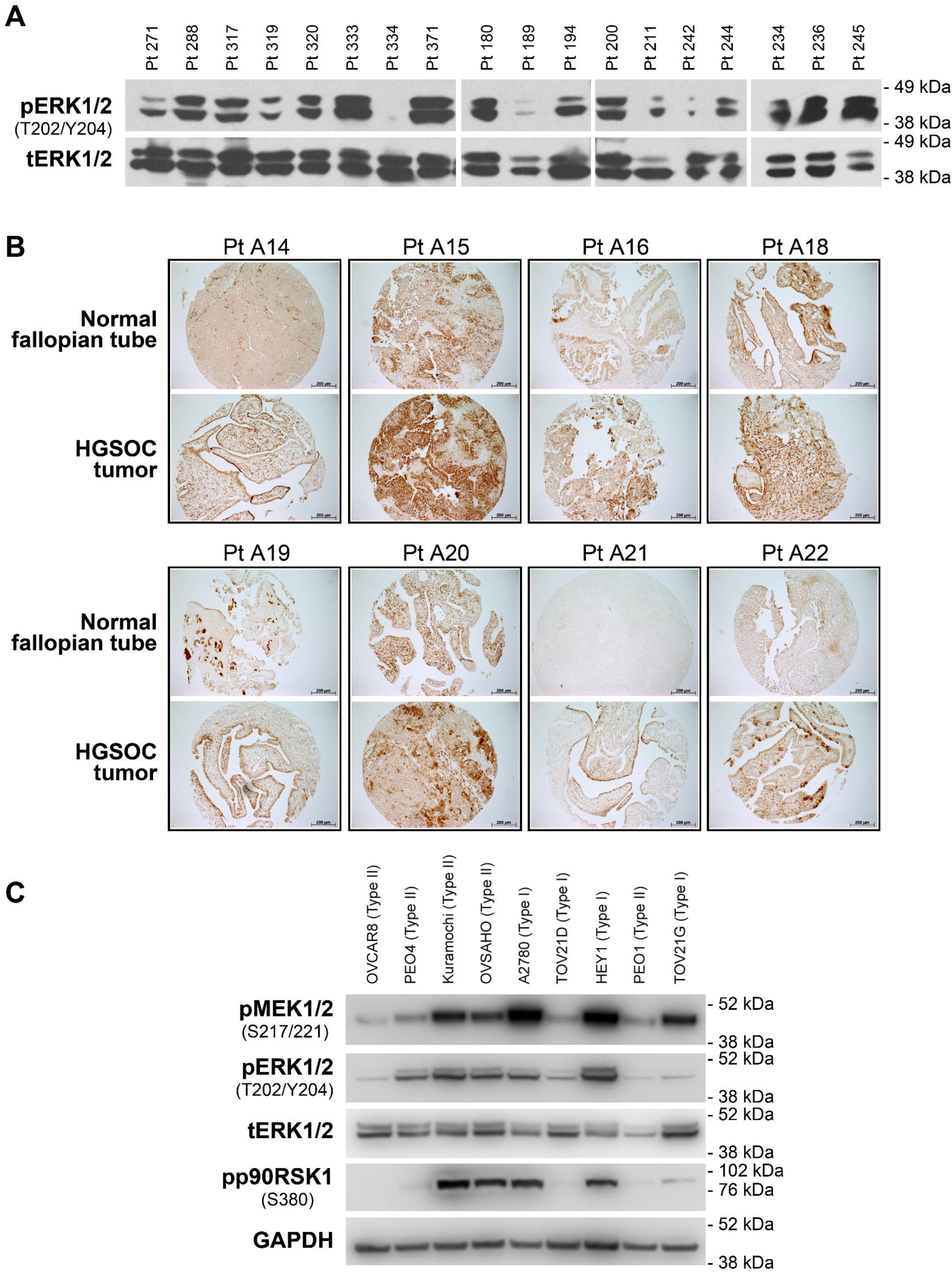
Hyperactivation of the MEK1/2 pathway in ovarian cancer tissue and cell lines. (A) Immunoblotting analysis of phosphorylated and total ERK1/2 levels in clinical samples of HGSOC tissue. (B) Immunohistochemical staining of phosphorylated ERK1/2 in paired clinical samples of HGSOC and normal fallopian tube tissues. (C) Immunoblotting analysis of MEK1/2-ERK1/2 pathway component activities in various ovarian cancer cell lines. pERK1/2 – phosphorylated ERK1/2, tERK1/2 – total ERK1/2, pMEK1/2 – phosphorylated MEK1/2, pp90RSK1 – phosphorylated p90RSK1, Pt XXX – patient number XXX.

The MEK1/2 pathway is also hyperactive in various ovarian cancer cell lines as indicated by high pMEK1/2, pERK1/2 and pp90RSK1 levels (Fig. 2C). Interestingly, treatment of OVCAR8 and PEO4 with cisplatin (Fig. S2) enhanced ERK1/2 and p90RSK1 activation in response to increasing doses of drug (Fig. 3A). We therefore generated cisplatin-resistant OVCAR8 and PEO4 cell cultures by prolonged cultivation of cells in the presence of 0.1 μg/mL or 0.25 μg/mL of cisplatin. In agreement with data reported above, development of cisplatin resistance was associated with enhanced ERK1/2 and p90RSK1 activity (Fig. 3B). To additionally confirm MEK1/2 signaling changes, we evaluated expression of 10 genes (*PHLDA1, SPRY2, SPRY4, DUSP4, DUSP6, CCND1, EPHA2, EPHA4, ETV4, ETV5*, see Table S3 for details) that were reported to reflect MEK1/2 activity [43]. We detected at least a two-fold increase in *PHLDA1, SPRY2, DUSP4, DUSP6*, and *EPHA2* expression in cisplatin-resistant cells (Fig. 3C). Together these results suggest a possible role of high MEK1/2-ERK1/2 activity in HGSOC development of chemoresistance.

**Fig. 3.**
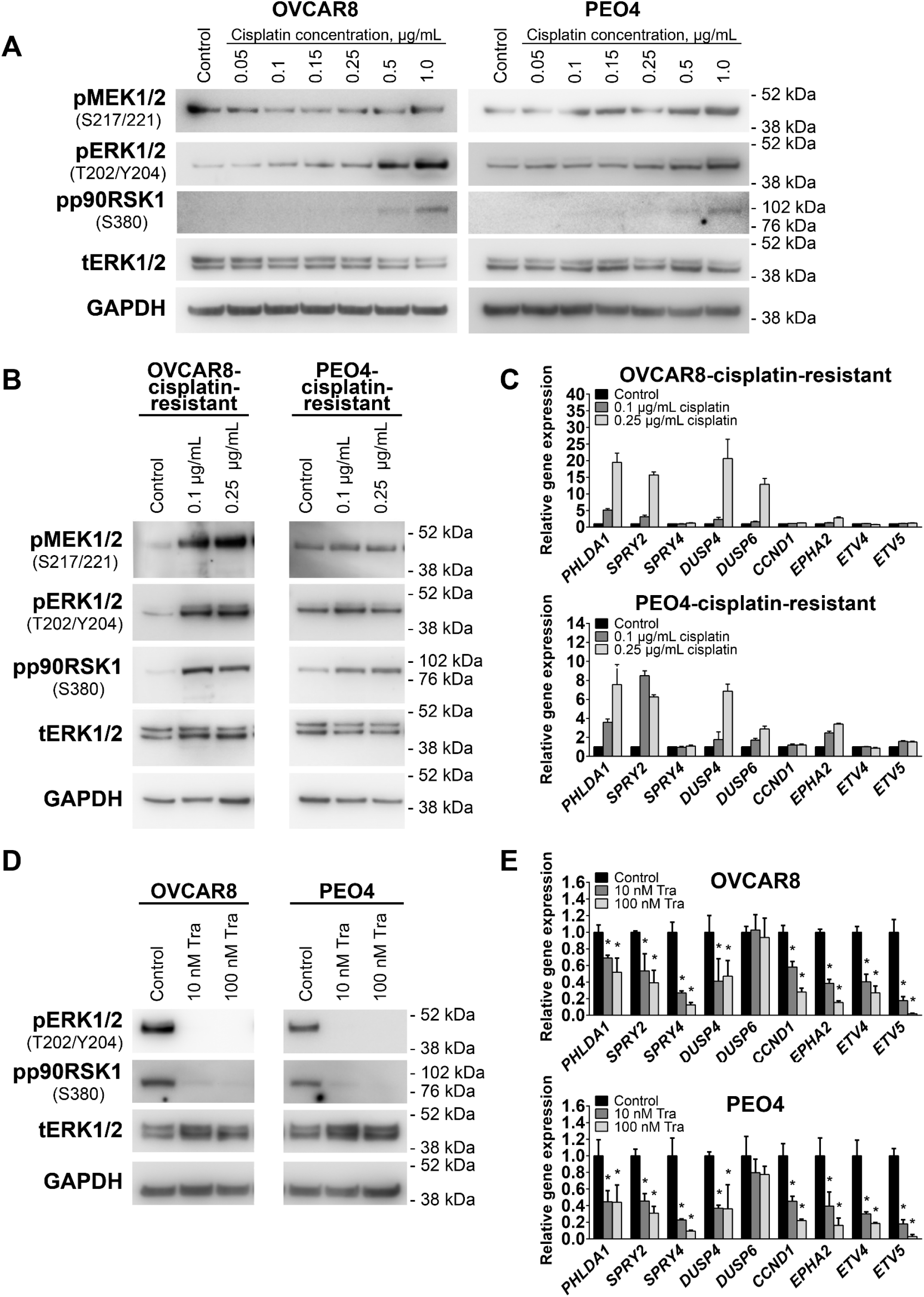
MEK1/2 pathway activity changes in HGSOC cell lines in response to drug treatment. (A) Immunoblotting analysis of MEK1/2-ERK1/2 pathway component activities in cells treated with vehicle or cisplatin for 24 hours. (B) Immunoblotting analysis of MEK1/2-ERK1/2 pathway component activities in cells resistant to the indicated concentrations of cisplatin. (C) Gene expression levels of MEK1/2-responsive genes in cells resistant to the indicated concentrations of cisplatin. Data are normalized to “Control” samples and presented as means ± S.E.M. (D) Immunoblotting analysis of MEK1/2-ERK1/2 pathway component activation in cells treated with vehicle or trametinib for 24 hours. (E) Gene expression levels of MEK1/2-responsive genes in cells treated with vehicle or trametinib for 10 hours. Data are normalized to “Control” samples and presented as means ± S.D. Tra – trametinib, pERK1/2 – phosphorylated ERK1/2, tERK1/2 – total ERK1/2, pMEK1/2 – phosphorylated MEK1/2, pp90RSK1 – phosphorylated p90RSK1.

### Inhibition of MEK1/2 causes arrest of HGSOC cell proliferation without inducing cell death

We next evaluated the impact of a selective and non-competitive FDA approved inhibitor MEK1/2 inhibitor, trametinib [36], on HGSOC proliferation. Activity of the MEK1/2 pathway was completely inhibited by 10 nM or higher concentrations of trametinib after 24 hours of treatment. Furthermore, trametinib-induced inactivation of the MEK1/2-ERK1/2 cascade caused corresponding decreases in the level of pp90RSK1, a downstream ERK1/2 target (Fig. 3D). MEK1/2 inhibition was also reflected in substantial dose-dependent downregulation of 8 MEK-responsive genes expressed in OVCAR8 and PEO4 cells after treatment with trametinib for 10 hours (Fig. 3E). These results indicate that trametinib is a potent inhibitor of MEK1/2 signaling activity in HGSOC cells that downregulates the activity of the entire MEK1/2-ERK1/2 axis, including downstream targets.

To evaluate the role of MEK1/2 activity on HGSOC proliferation, we treated cells for 72 hours with a wide range of trametinib concentrations (0.5 nM – 1000 nM). The resulting IC_50_ and GI_50_ values were 8.4 nM and 10.2 nM for OVCAR8 cells and 5.5 nM and 6.5 nM for PEO4 cells, respectively (Fig. 4A), with a maximum effect at 100 nM. Based on these results, two trametinib concentrations (10 nM and 100 nM) were chosen for further experiments. Treatment with 100 nM trametinib reduced viable cell numbers to 15% (OVCAR8) and 13% (PEO4) of control values (Fig. 4B) without causing considerable cytotoxic action (Fig. 4C, S3). Cisplatin-resistant OVCAR8 and PEO4 cells displayed higher sensitivity to cytostatic but not cytotoxic effects of trametinib treatment (Fig. S4). Imaging of live cells treated with trametinib revealed an increase in cell size (Fig. 4D). Cell cycle analysis demonstrated that trametinib treatment caused arrest of proliferation in the G_1/0_-phase but did not affect the sub-G_0_ fraction of apoptotic cells (Fig. 4E).

**Fig. 4.**
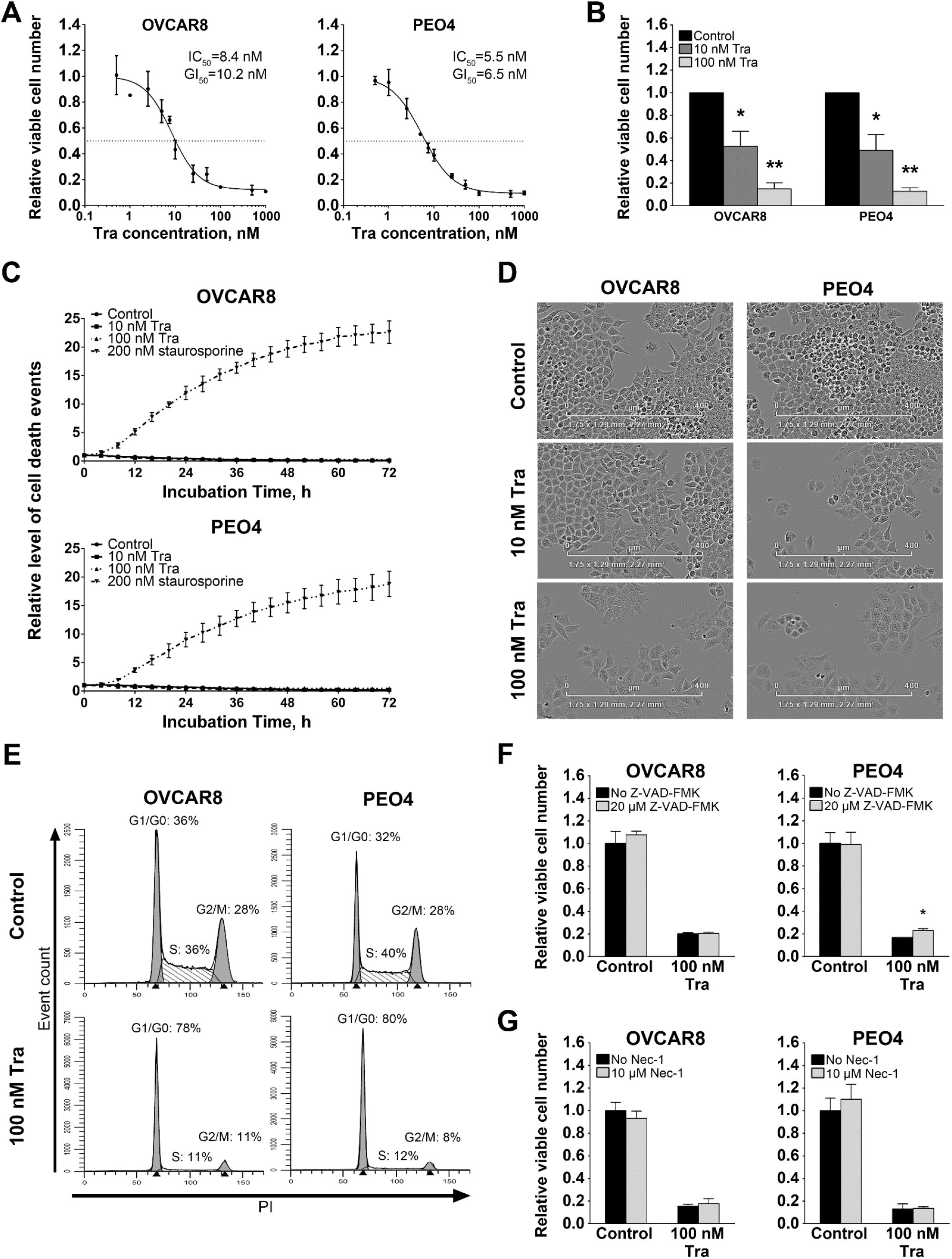
Changes in functional characteristics of HGSOC cell cultures caused by trametinib treatment. (A) Dose-response curves generated using relative viable cell numbers after treatment with various concentrations of trametinib for 72 hours. Data are normalized to vehicle-treated control samples (not shown) and presented as means ± S.D. (B) Viable cell numbers after treatment with selected concentrations of trametinib for 72 hours. Data are normalized to “Control” samples and presented as means ± S.D. (C) Cytotoxic effect of trametinib treatment detected using the CytoTox reagent. Staurosporin is used as a positive control. Fluorescence level for each time point is normalized to the area covered by cells and starting value; data are presented as means ± S.D. (D) Cell morphology and confluence after treatment with trametinib for 72 hours. (E) Cell cycle phase analysis after trametinib treatment for 24 hours. (F) Effect of pan-caspase inhibitor Z-VAD-FMK on cells treated with vehicle or trametinib for 72 hours. Data are normalized to “Control, No Z-VAD-FMK” sample and presented as means ± S.D. (G) Effect of the RIPK1 inhibitor necrostatin-1 on cells treated with vehicle or trametinib for 72 hours. Data are normalized to “Control, No Nec-1” sample and presented as means ± S.D. Tra – trametinib, PI – propidium iodide, Nec-1 – necrostatin-1.

While caspase inhibition with Z-VAD-FMK attenuated apoptosis induction by 0.2 μM staurosporine (positive control, Fig. S5A), it did not impact trametinib-induced changes in viable cell numbers (Fig. 4F). Similarly, inhibition of the major necroptosis regulator RIPK1 with 10 μM Nec-1 (Fig. 4G) or siRNA-mediated knockdown of *RIPK1* expression (Fig. S5B, S5C) did not affect trametinib-treated cells, suggesting that the reduction in cell number is not due to either caspase-dependent apoptosis or RIPK1-dependent necroptosis.

### Trametinib promotes stemness of HGSOC cells

Because CSCs are associated with chemoresistance and recurrence [18, 19, 23, 24], we investigated the effect of trametinib on cancer stemness. Both OVCAR8 and PEO4 cell lines display high percentages of CD133^+^ cells that are unaffected by trametinib treatment (Fig. 5A). In contrast, trametinib treatment of PEO4 cells significantly promoted the expression of cell stemness regulators *SOX2, NANOG, POU5F1* (*OCT4*), and two *ALDH1A* homologs associated with elevated chemoresistance and the stem-like phenotype of ovarian cancer cells [19, 23, 44, 45] (Fig. 5B). We next treated PEO4 cells with vehicle or 100 nM trametinib for 72 hours, then performed drug washout to propagate the cells surviving the treatment (hereinafter denoted as “PEO4-Washout-Control” and “PEO4-Washout-Tra”, Fig. S6A). PEO4-Washout-Tra cells displayed no significant differences in activity of MEK1/2 or other signaling pathways (Fig. 5C) or expression levels of MEK1/2-responsive genes (Fig. S6B) in comparison to control cells. Trametinib-driven selection had no biologically relevant effect upon cell proliferation rate (Fig. S6C), sensitivity to cisplatin (Fig. S6D), or percentage of CD133^+^ cells (Fig. S6E); but resulted in *SOX2* and *ALDHA1A* expression upregulation in comparison to control cells (Fig. 5D). PEO4-Washout-Tra also displayed enrichment of cells with very high ALDH activity (Fig 5E, “High” gate).

**Fig. 5.**
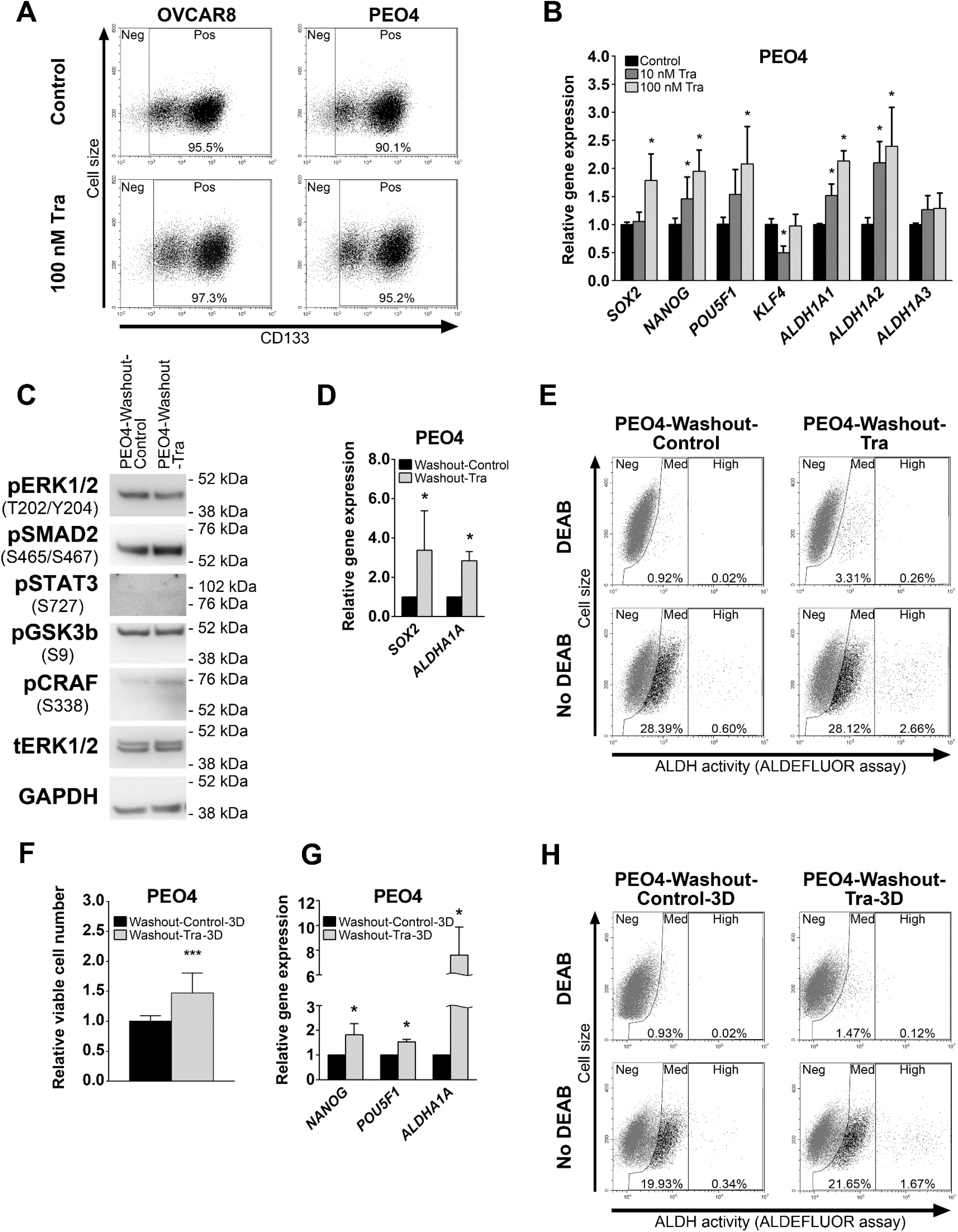
Effects of trametinib treatment on stemness-related characteristics of HGSOC cells. (A) Expression of the CD133 surface marker in cells treated with vehicle and trametinib for 72 hours. Gates indicate CD133-negative (“Neg”) and CD133-positive (“Pos”) cell subpopulations. (B) Gene expression levels of stemness-related genes in cells treated with vehicle or trametinib for 10 hours. Data are normalized to “Control” samples and presented as means ± S.D. (C) Immunoblotting analysis of various signaling proteins activation in PEO4-Washout cells. (D) Gene expression levels of stemness-related genes in PEO4-Washout cells. Data are normalized to “Washout-Control” samples and presented as means ± S.D. (E) ALDEFLUOR analysis of PEO4-Washout cells. Gates indicate cell subpopulations displaying negative (“Neg”), medium (“Med”), or high (“High”) levels of ALDH activity. (F) Viable cell numbers of PEO4-Washout cells after cultivation in non-adherent 3D conditions for 7 days. Data are normalized to “Washout-Control-3D” samples and presented as means ± S.D. (G) Gene expression levels of stemness-related genes in PEO4-Washout cells after cultivation in non-adherent 3D conditions for 7 days. Data are normalized to “Washout-Control-3D” samples and presented as means ± S.D. (H) ALDEFLUOR analysis of PEO4-Washout cells after cultivation in non-adherent 3D conditions for 7 days. Gates indicate cell subpopulations displaying negative (“Neg”), medium (“Med”), or high (“High”) levels of ALDH activity.

The ability to grow under non-adherent conditions is a distinct feature of CSCs. Thus we conducted a spheroid formation assay and detected a significantly higher growth rate of PEO4-Washout-Tra cells in comparison to control (Fig. 5F). Moreover, induction of *ALDH1A1* expression observed in adherent PEO4-Washout-Tra cells was further increased in spheroids and was accompanied by upregulation of *NANOG* and *POU5F1* expression (Fig. 5G). PEO4-Washout-Tra cells grown as spheroids retained a very high percentage of CD133^+^ cells (> 90%, Fig. S6F) and a higher fraction of ALDEFLUOR-bright cells compared to control cells (Fig. 5H, “High” gate). Taken together, these results indicate that cells surviving trametinib treatment obtain a more pronounced CSC phenotype.

### *Effect of MEK1/2 inhibition* in vivo

To examine the impact of MEK1/2 inhibition *in vivo*, we injected PEO4 cells subcutaneously into mice that were subsequently treated intraperitoneally with 1 mg/kg trametinib or vehicle daily. Trametinib treatment significantly reduced the rate of tumor growth and caused a 4-fold decrease in tumor volume as estimated at 4 weeks after initial cell injection (Fig 6A). Tumors grown in mice from both control and trametinib-treated experimental groups displayed tissue morphology typical of HGSOC (Fig. 6B). Immunoblotting analysis of tumor tissue samples demonstrated that trametinib treatment predominantly inhibited phosphorylation of ERK1 (Fig. 6C, upper band detected by pERK1/2 antibodies). Inhibition of MEK1/2-ERK1/2 activity was confirmed by RT-qPCR analysis that revealed trametinib induced decreases in *PHLDA1, SPRY4, DUSP4, DUSP6, EPHA2, ETV4*, and *ETV5* expression (Fig. 6D). In contrast with the changes observed in cells, *SPRY2* and *CCND1* expression was not affected by trametinib *in vivo*. Thus, trametinib treatment of tumors *in vivo* caused prominent inhibition of MEK1/2 pathway activity and the rate of tumor growth in full concordance with results obtained in cells.

**Fig. 6.**
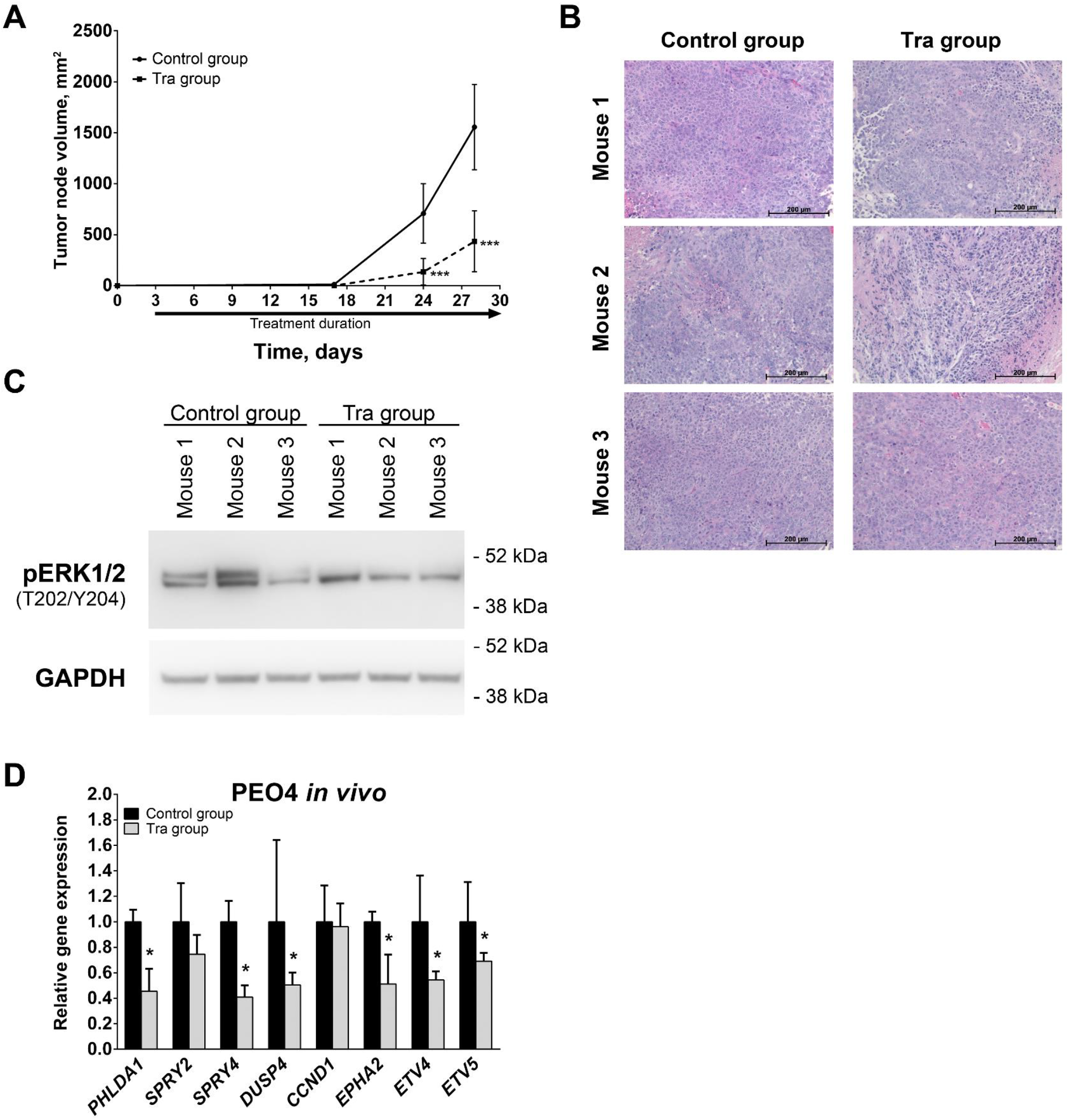
Effects of trametinib treatment on HGSOC growth *in vivo*. (A) Growth kinetics of tumors developed from subcutaneously injected PEO4 cells. (B) Hematoxylin and eosin staining of PEO4 xenograft tissue samples. (C) Immunoblotting analysis of phosphorylated ERK1/2 levels in PEO4 xenograft tissue samples. (D) Gene expression levels of MEK1/2-responsive genes in PEO4 xenograft tissue samples. Data are normalized to “Control group” samples and presented as means ± S.D. Tra – trametinib, pERK1/2 – phosphorylated ERK1/2.

## Discussion

The present study evaluated the role of the MEK1/2 pathway in HGSOC and assessed MEK1/2 inhibition as a therapeutic approach. Activation of the MAPK signaling pathway is one of the most frequent events in cancer and affects many features inherent to malignant cells [32, 46]. Most notably, high activity of the MEK1/2-ERK1/2 portion of the MAPK cascade directly promotes cell proliferation, survival, and drug resistance [31–33, 47]. Furthermore, a number of studies reported that ERK1/2 activation occurs in CSCs [27] and is crucial for cell survival and proliferation in prostate, breast, and thyroid tumors [28–30]. A special focus in MAPK signaling inhibition is placed on MEK1/2 as it acts as a “gatekeeper” of MAPK pathway, conducting the signal from multiple upstream regulators towards ERK1/2, which are the only downstream targets of MEK1/2 [34]. Development of a new generation of non-competitive, highly specific MEK1/2 inhibitors (trametinib [48], selumetinib [49], cobimetinib [50], and others) that show high efficiency and tolerable side effects significantly increases the therapeutic potential of MEK1/2 inhibition.

Despite a lack of mutations activating *KRAS* or *BRAF* in HGSOCs, we observed high levels of MEK1/2-ERK1/2 activity in a majority of clinical samples and cell lines. This observation is further supported by recently published studies reporting an association between high MAPK activity in HGSOC and poor survival [39, 51]. Moreover, cisplatin treatment of HGSOC cells results in further MEK1/2 pathway activation that persists in cisplatin-resistant cells. MEK1/2-ERK1/2 signaling inhibition in ovarian cancer cells is reported to sensitize them to chemotherapy [47], so MEK1/2 pathway hyperactivation could be a mechanism allowing HGSOC cells to overcome cytotoxic effects of cisplatin. Cisplatin-induced MEK1/2 activation in OVCAR8 and PEO4 cell lines established from recurrent tumors, which already obtained cisplatin resistance [52–54], suggests that repeated cisplatin treatment can further promote MEK1/2-ERK1/2 activity and associated chemoresistance in HGSOC. Such an effect could possibly be facilitated through cisplatin-driven enrichment of chemoresistant CSCs that often display high MEK1/2 pathway activity.

TCGA data suggest that MEK1/2 signaling hyperactivation is caused by genetic amplifications and overexpression of upstream MAPK components (*GRB2, SOS1, RAS*, and *RAF* families) and therefore could be countered by MEK1/2 inhibition. Indeed, treatment of ovarian cancer cell lines with the MEK1/2 inhibitor, selumetinib, results in prominent inhibition of ERK1/2 phosphorylation [51]. Considering the MEK1/2-activating effects of cisplatin discussed above, we focused our studies on cisplatin-resistant cells, OVCAR8 and PEO4. These cell lines also display high ALDH1 activity and prominent subpopulations of CD133^+^ cells, two distinct characteristics of ovarian CSCs [23, 44, 55–58].

Trametinib treatment drastically reduces the growth rates of HGSOC cells both in culture and *in vivo*, confirming that MEK1/2-ERK1/2 activity is involved in driving cell proliferation [31]. Anti-proliferative effects of MEK1/2 inhibitors have been reported in many cell types, including ovarian cancer cells, and cause G_1/0_-phase cell cycle arrest due to loss of ERK1/2 activation [39, 51, 59, 60]. However, despite pronounced cytostatic action *in vitro*, most first generation MEK1/2 inhibitors demonstrated limited efficacy in clinical trials involving melanoma, breast, colon, lung, and pancreatic cancer patients [35, 61]. Trametinib, on the other hand, belongs to a new generation of MEK1/2 inhibitors, displays higher efficiency, and was the first MEK1/2 inhibitor approved by FDA for cancer treatment [36]. Thus, trametinib could possibly show greater efficacy in clinical trials.

We observed that trametinib induces cell cycle arrest in HGSOC lines. The potential of cell cycle arrest in HGSOC is best illustrated by paclitaxel that blocks cell cycle in the G2/M-phase. Prolonged mitotic block results in apoptosis induction and eventual cell death [62, 63]. In a similar way, inhibition of MEK1/2 with trametinib induced death of various tumor cells that are heavily dependent on elevated RAS-RAF-MEK1/2-ERK1/2 cascade activity [64–67]. In contrast to these reports, we did not detect any cytotoxic effects caused by MEK1/2 inhibition in HGSOC cells. Recently another MEK1/2 inhibitor, selumetinib, was reported to induce apoptosis in the PEO1 HGSOC cell line [51]. This discrepancy can be explained by the fact that both OVCAR8 and PEO4 cells represent cisplatin-resistant subgroups of HGSOC in comparison to cisplatin-sensitive PEO1 cells and therefore may have developed various means of avoiding cell death. Our results indicate that active MEK1/2 signaling, while promoting proliferation of cisplatin-resistant HGSOC cells, is not essential for their survival. Moreover, loss of MEK1/2-regulated negative feedback can induce RAF hyperactivation that promotes cell viability through MEK1/2-unrelated pathways [61].

Cells surviving trametinib treatment obtain a prominent stem-like phenotype, including increased ability to grow under non-adherent conditions, increased ALDH1 activity, and a very high CD133^+^ fraction typical of ovarian CSCs [58]. This effect could be due to trametinib favoring the propagation of pre-existing CSC subpopulations or directly inducing stem-like properties in affected cells. Our results indicate that the latter option is more likely as trametinib increased expression of stemness-related genes after only 10 hours of treatment. Furthermore, trametinib-driven cell stemness was persistent for at least 10 passages after drug washout despite MEK1/2-ERK1/2 activity being restored to control levels. Described effects are in disagreement with the previously reported anti-stemness impact of MEK1/2 inhibition in PEO1 cells [51], suggesting differences between cisplatin-sensitive and cisplatin-resistant HGSOC cells. Consistent with our findings, ALDH1 overexpression in response to BRAF and MEK1/2 inhibition was recently reported in melanoma [68]. Given that high expression of ALDH1 and CD133 in ovarian tumors is strongly associated with poor prognosis and chemoresistance [69–73], we conclude that prolonged treatment of cisplatin-resistant HGSOC with trametinib might promote CSC enrichment.

The present study demonstrates that treatment with trametinib as a single drug delays HGSOC tumor growth and therefore has the potential for prolonging disease-free survival of HGSOC patients. Because cisplatin-resistant cells are more sensitive to trametinib, this drug may prove to be an important therapeutic option for platinum-resistant recurrent tumors and initially cisplatin-refractory cases. Due to its fewer side effects, trametinib treatment can also be considered for patients who are unable to tolerate certain chemotherapeutic regimens due to systemic toxicity effects. Because trametinib treatments are associated with cancer cell stemness, its combination with other targeted therapies showing higher cytotoxic effects and, ideally, targeting ovarian CSCs may be key in treatment of HGSOC. Because cells surviving trametinib treatment display very high ALDH1A expression and activity, combination with ALDH inhibitors may also offer benefit. This suggestion is further supported by increases in nifuroxazide sensitivity in melanoma cells overexpressing ALDH1 due to MEK1/2 inhibition [68]. Recently a selective inhibitor of the ALDH1A family was identified as a chemical agent capable of efficiently inducing necroptotic death in ovarian CSCs [23]. The combination of trametinib and an ALDH1A inhibitor could retain the tumor growth arresting effect of trametinib while further complementing it by eliminating the surviving ALDH+ tumor cells in a targeted manner.

## Acknowledgments

This work was supported in part by the Michigan Ovarian Cancer Alliance (MIOCA) and The Foundation for Women’s Cancer (FWC) to I.C. R.B is supported by NIH NCI 1R01CA218026.

## Supplementary Figure Legends

**Fig. S1.** Hematoxylin and eosin staining of paired clinical samples of HGSOC and normal fallopian tube tissues. Pt XXX – patient number XXX.

**Fig. S2.** Dose-response curves generated using relative viable cell numbers of HGSOC cells after treatment with various concentrations of cisplatin for 72 hours. Data are normalized to vehicle-treated control samples (not shown) and presented as means ± S.D.

**Fig. S3.** Viable cell percentage of HGSOC cells after treatment with selected concentrations of trametinib for 72 hours. Data are normalized to “Control” samples and presented as means ± S.D.

**Fig. S4.** Viable cell numbers (A) and viable cell percentages (B) of cisplatin-resistant OVCAR8 and PEO4 cells after treatment with selected concentrations of trametinib for 72 hours. Data for viable cell numbers are normalized to “Control”-”No Tra” samples for each cell culture and presented as mean+SD.

**Fig. S5. Effect of cell death-related treatment of HGSOC cells.** (A) Effect of the pan-caspase inhibitor Z-VAD-FMK on cells treated with vehicle or staurosporine for 24 hours. Data are normalized to “Control, No Z-VAD-FMK” sample and presented as means ± S.D. (B) Gene expression levels of the *RIPK1* gene in HGSOC cells transfected with anti-RIPK1 siRNA. Data are normalized to “No Target siRNA” samples and presented as means ± S.E.M. (C) Effect of siRNA-mediated *RIPK1* knockdown upon cells treated with vehicle or trametinib for 72 hours. Data are normalized to “Control, No Target siRNA” sample and presented as means ± S.D. Tra – trametinib, siRNA – short interfering RNA.

**Fig. S6. Effects of trametinib washout upon properties of PEO4 cells.** (A) Growth kinetics of PEO4 cells during the initial establishment of PEO4-Washout cells. Data are normalized to starting cell number and presented as means ± S.D. (B) Gene expression levels of MEK1/2-responsive genes in PEO4-Washout cells. Data are normalized to “Washout-Control” samples and presented as means ± S.E.M. (C) Growth kinetics of PEO4-Washout cells in standard conditions. Data are normalized to starting cell number and presented as means ± S.D. (D) Dose-response curves generated using relative viable numbers of PEO4-Washout cells after treatment with various concentrations of cisplatin for 72 hours. Data are normalized to vehicle-treated control samples (not shown) and presented as means ± SD. (E) Expression of the CD133 surface marker in PEO4-Washout cells grown in standard conditions. Gates indicate CD133-negative (“Neg”) and CD133-positive (“Pos”) cell subpopulations. (F) Expression of the CD133 surface marker in PEO4-Washout cells grown in non-adherent conditions. Gates indicate CD133-negative (“Neg”) and CD133-positive (“Pos”) cell subpopulations. Tra – trametinib.

## References

1. Siegel RL, Miller KD, Jemal A. Cancer Statistics, 2017. CA: a cancer journal for clinicians. 2017;67(1):7–30. Epub 2017/01/06. doi: 10.3322/caac.21387. PubMed PMID: 28055103.

2. Torre LA, Trabert B, DeSantis CE, Miller KD, Samimi G, Runowicz CD, et al. Ovarian cancer statistics, 2018. CA: a cancer journal for clinicians. 2018;68(4):284–96. Epub 2018/05/29. doi: 10.3322/caac.21456. PubMed PMID: 29809280.

3. Kurman RJ, Shih Ie M. The Dualistic Model of Ovarian Carcinogenesis: Revisited, Revised, and Expanded. The American journal of pathology. 2016;186(4):733–47. Epub 2016/03/26. doi: 10.1016/j.ajpath.2015.11.011. PubMed PMID: 27012190; PubMed Central PMCID: PMCPMC5808151.

4. Cooke SL, Brenton JD. Evolution of platinum resistance in high-grade serous ovarian cancer. The Lancet Oncology. 2011;12(12):1169–74. Epub 2011/07/12. doi: 10.1016/S1470-2045(11)70123-1. PubMed PMID: 21742554.

5. Vaughan S, Coward JI, Bast RC, Jr., Berchuck A, Berek JS, Brenton JD, et al. Rethinking ovarian cancer: recommendations for improving outcomes. Nat Rev Cancer. 2011;11(10):719–25. Epub 2011/09/24. doi: 10.1038/nrc3144. PubMed PMID: 21941283; PubMed Central PMCID: PMCPMC3380637.

6. The Surveillance, Epidemiology, and End Results (SEER) Program of the National Cancer Institute (NCI) [11.08.18]. Available from: https://seer.cancer.gov/statfacts/html/ovary.html.

7. Morgan RJ, Jr., Armstrong DK, Alvarez RD, Bakkum-Gamez JN, Behbakht K, Chen LM, et al. Ovarian Cancer, Version 1.2016, NCCN Clinical Practice Guidelines in Oncology. Journal of the National Comprehensive Cancer Network: JNCCN. 2016;14(9):1134–63. Epub 2016/09/03. PubMed PMID: 27587625.

8. Raja FA, Chopra N, Ledermann JA. Optimal first-line treatment in ovarian cancer. Annals of oncology: official journal of the European Society for Medical Oncology. 2012; 23 Suppl 10:x118-27. Epub 2012/09/26. doi: 10.1093/annonc/mds315. PubMed PMID: 22987945.

9. Davis A, Tinker AV, Friedlander M. “Platinum resistant” ovarian cancer: what is it, who to treat and how to measure benefit? Gynecologic oncology. 2014;133(3):624–31. Epub 2014/03/13. doi: 10.1016/j.ygyno.2014.02.038. PubMed PMID: 24607285.

10. Cortez AJ, Tudrej P, Kujawa KA, Lisowska KM. Advances in ovarian cancer therapy. Cancer chemotherapy and pharmacology. 2018;81(1):17–38. Epub 2017/12/19. doi: 10.1007/s00280-017-3501-8. PubMed PMID: 29249039; PubMed Central PMCID: PMCPMC5754410.

11. Christie EL, Bowtell DDL. Acquired chemotherapy resistance in ovarian cancer. Annals of oncology: official journal of the European Society for Medical Oncology. 2017;28(suppl_8): viii13–viii5. Epub 2017/12/13. doi: 10.1093/annonc/mdx446. PubMed PMID: 29232469.

12. Agarwal R, Kaye SB. Ovarian cancer: strategies for overcoming resistance to chemotherapy. Nature reviews Cancer. 2003;3(7):502–16. Epub 2003/07/02. doi: 10.1038/nrc1123. PubMed PMID: 12835670.

13. Freimund AE, Beach JA, Christie EL, Bowtell DDL. Mechanisms of Drug Resistance in High-Grade Serous Ovarian Cancer. Hematology/oncology clinics of North America. 2018;32(6):983–96. Epub 2018/11/06. doi: 10.1016/j.hoc.2018.07.007. PubMed PMID: 30390769.

14. Norouzi-Barough L, Sarookhani MR, Sharifi M, Moghbelinejad S, Jangjoo S, Salehi R. Molecular mechanisms of drug resistance in ovarian cancer. Journal of cellular physiology. 2018;233(6):4546–62. Epub 2017/11/21. doi: 10.1002/jcp.26289. PubMed PMID: 29152737.

15. Mittempergher L. Genomic Characterization of High-Grade Serous Ovarian Cancer: Dissecting Its Molecular Heterogeneity as a Road Towards Effective Therapeutic Strategies. Current oncology reports. 2016;18(7):44. Epub 2016/06/01. doi: 10.1007/s11912-016-0526-9. PubMed PMID: 27241520.

16. Yin X, Jing Y, Cai MC, Ma P, Zhang Y, Xu C, et al. Clonality, Heterogeneity, and Evolution of Synchronous Bilateral Ovarian Cancer. Cancer research. 2017;77(23):6551–61. Epub 2017/10/04. doi: 10.1158/0008-5472.CAN-17-1461. PubMed PMID: 28972072.

17. Cooke SL, Ng CK, Melnyk N, Garcia MJ, Hardcastle T, Temple J, et al. Genomic analysis of genetic heterogeneity and evolution in high-grade serous ovarian carcinoma. Oncogene. 2010;29(35):4905–13. Epub 2010/06/29. doi: 10.1038/onc.2010.245. PubMed PMID: 20581869; PubMed Central PMCID: PMCPMC2933510.

18. Chefetz I, Alvero AB, Holmberg JC, Lebowitz N, Craveiro V, Yang-Hartwich Y, et al. TLR2 enhances ovarian cancer stem cell self-renewal and promotes tumor repair and recurrence. Cell Cycle. 2013;12(3):511–21. Epub 2013/01/18. doi: 10.4161/cc.23406 23406 [pii]. PubMed PMID: 23324344; PubMed Central PMCID: PMC3587452.

19. Mor G, Yin G, Chefetz I, Yang Y, Alvero A. Ovarian cancer stem cells and inflammation. Cancer biology & therapy. 2011;11(8):708–13. Epub 2011/02/15. PubMed PMID: 21317559; PubMed Central PMCID: PMCPMC3100563.

20. Zhang QH, Dou HT, Xu P, Zhuang SC, Liu PS. Tumor recurrence and drug resistance properties of side population cells in high grade ovary cancer. Drug research. 2015;65(3):153–7. Epub 2014/12/17. doi: 10.1055/s-0034-1375609. PubMed PMID: 25504004.

21. Kuroda T, Hirohashi Y, Torigoe T, Yasuda K, Takahashi A, Asanuma H, et al. ALDH1-high ovarian cancer stem-like cells can be isolated from serous and clear cell adenocarcinoma cells, and ALDH1 high expression is associated with poor prognosis. PloS one. 2013;8(6):e65158. Epub 2013/06/14. doi: 10.1371/journal.pone.0065158. PubMed PMID: 23762304; PubMed Central PMCID: PMCPMC3675199.

22. Ruscito I, Cacsire Castillo-Tong D, Vergote I, Ignat I, Stanske M, Vanderstichele A, et al. Exploring the clonal evolution of CD133/aldehyde-dehydrogenase-1 (ALDH1)-positive cancer stem-like cells from primary to recurrent high-grade serous ovarian cancer (HGSOC). A study of the Ovarian Cancer Therapy-Innovative Models Prolong Survival (OCTIPS) Consortium. European journal of cancer (Oxford, England: 1990). 2017;79:214–25. Epub 2017/05/20. doi: 10.1016/j.ejca.2017.04.016. PubMed PMID: 28525846.

23. Chefetz I, Grimley E, Yang K, Hong L, Vinogradova EV, Suciu R, et al. A Pan-ALDH1A Inhibitor Induces Necroptosis in Ovarian Cancer Stem-like Cells. Cell reports. 2019;26(11):3061–75.e6. Epub 2019/03/14. doi: 10.1016/j.celrep.2019.02.032. PubMed PMID: 30865894.

24. Chefetz I, Holmberg JC, Alvero AB, Visintin I, Mor G. Inhibition of Aurora-A kinase induces cell cycle arrest in epithelial ovarian cancer stem cells by affecting NFkB pathway. Cell cycle (Georgetown, Tex). 2011;10(13):2206–14. Epub 2011/05/31. doi: 10.4161/cc.10.13.16348. PubMed PMID: 21623171; PubMed Central PMCID: PMCPMC3154367.

25. O’Connor ML, Xiang D, Shigdar S, Macdonald J, Li Y, Wang T, et al. Cancer stem cells: A contentious hypothesis now moving forward. Cancer letters. 2014;344(2):180–7. Epub 2013/12/18. doi: 10.1016/j.canlet.2013.11.012. PubMed PMID: 24333726.

26. Dean M, Fojo T, Bates S. Tumour stem cells and drug resistance. Nature reviews Cancer. 2005;5(4):275–84. Epub 2005/04/02. doi: 10.1038/nrc1590. PubMed PMID: 15803154.

27. Martins-Neves SR, Cleton-Jansen AM, Gomes CMF. Therapy-induced enrichment of cancer stem-like cells in solid human tumors: Where do we stand? Pharmacological research. 2018;137:193–204. Epub 2018/10/15. doi: 10.1016/j.phrs.2018.10.011. PubMed PMID: 30316903.

28. Rybak AP, Ingram AJ, Tang D. Propagation of human prostate cancer stem-like cells occurs through EGFR-mediated ERK activation. PloS one. 2013;8(4):e61716. Epub 2013/04/27. doi: 10.1371/journal.pone.0061716. PubMed PMID: 23620784; PubMed Central PMCID: PMCPMC3631151.

29. Ahn HJ, Kim G, Park KS. Ell3 stimulates proliferation, drug resistance, and cancer stem cell properties of breast cancer cells via a MEK/ERK-dependent signaling pathway. Biochemical and biophysical research communications. 2013;437(4):557–64. Epub 2013/07/16. doi: 10.1016/j.bbrc.2013.06.114. PubMed PMID: 23850691.

30. Wang Y, Lin X, Fu X, Yan W, Lin F, Kuang P, et al. Long non-coding RNA BANCR regulates cancer stem cell markers in papillary thyroid cancer via the RAF/MEK/ERK signaling pathway. Oncology reports. 2018;40(2):859–66. Epub 2018/06/20. doi: 10.3892/or.2018.6502. PubMed PMID: 29917164.

31. Zhang W, Liu HT. MAPK signal pathways in the regulation of cell proliferation in mammalian cells. Cell research. 2002;12(1):9–18. Epub 2002/04/11. doi: 10.1038/sj.cr.7290105. PubMed PMID: 11942415.

32. Burotto M, Chiou VL, Lee JM, Kohn EC. The MAPK pathway across different malignancies: a new perspective. Cancer. 2014;120(22):3446–56. Epub 2014/06/21. doi: 10.1002/cncr.28864. PubMed PMID: 24948110; PubMed Central PMCID: PMCPMC4221543.

33. McCubrey JA, Steelman LS, Chappell WH, Abrams SL, Wong EW, Chang F, et al. Roles of the Raf/MEK/ERK pathway in cell growth, malignant transformation and drug resistance. Biochimica et biophysica acta. 2007;1773(8):1263–84. Epub 2006/11/28. doi: 10.1016/j.bbamcr.2006.10.001. PubMed PMID: 17126425; PubMed Central PMCID: PMCPMC2696318.

34. Caunt CJ, Sale MJ, Smith PD, Cook SJ. MEK1 and MEK2 inhibitors and cancer therapy: the long and winding road. Nature reviews Cancer. 2015;15(10):577–92. Epub 2015/09/25. doi: 10.1038/nrc4000. PubMed PMID: 26399658.

35. Mandal R, Becker S, Strebhardt K. Stamping out RAF and MEK1/2 to inhibit the ERK1/2 pathway: an emerging threat to anticancer therapy. Oncogene. 2016;35(20):2547–61. Epub 2015/09/15. doi: 10.1038/onc.2015.329. PubMed PMID: 26364606.

36. U.S. Food and Drug Administration [11.08.18]. Available from: https://www.fda.gov/.

37. Singer G, Oldt R, 3rd, Cohen Y, Wang BG, Sidransky D, Kurman RJ, et al. Mutations in BRAF and KRAS characterize the development of low-grade ovarian serous carcinoma. Journal of the National Cancer Institute. 2003;95(6):484–6. Epub 2003/03/20. PubMed PMID: 12644542.

38. Miller CR, Oliver KE, Farley JH. MEK1/2 inhibitors in the treatment of gynecologic malignancies. Gynecologic oncology. 2014;133(1):128–37. Epub 2014/01/18. doi: 10.1016/j.ygyno.2014.01.008. PubMed PMID: 24434059.

39. Hew KE, Miller PC, El-Ashry D, Sun J, Besser AH, Ince TA, et al. MAPK Activation Predicts Poor Outcome and the MEK Inhibitor, Selumetinib, Reverses Antiestrogen Resistance in ER-Positive High-Grade Serous Ovarian Cancer. Clinical cancer research: an official journal of the American Association for Cancer Research. 2016;22(4):935–47. Epub 2015/10/21. doi: 10.1158/1078-0432.ccr-15-0534. PubMed PMID: 26482043; PubMed Central PMCID: PMCPMC4755805.

40. Pfaffl MW. A new mathematical model for relative quantification in real-time RT-PCR. Nucleic acids research. 2001;29(9):e45. Epub 2001/05/09. PubMed PMID: 11328886; PubMed Central PMCID: PMCPMC55695.

41. Gao J, Aksoy BA, Dogrusoz U, Dresdner G, Gross B, Sumer SO, et al. Integrative analysis of complex cancer genomics and clinical profiles using the cBioPortal. Science signaling. 2013;6(269):pl1. Epub 2013/04/04. doi: 10.1126/scisignal.2004088. PubMed PMID: 23550210; PubMed Central PMCID: PMCPMC4160307.

42. Cerami E, Gao J, Dogrusoz U, Gross BE, Sumer SO, Aksoy BA, et al. The cBio cancer genomics portal: an open platform for exploring multidimensional cancer genomics data. Cancer discovery. 2012;2(5):401–4. Epub 2012/05/17. doi: 10.1158/2159-8290.cd-12-0095. PubMed PMID: 22588877; PubMed Central PMCID: PMCPMC3956037.

43. Wagle MC, Kirouac D, Klijn C, Liu B, Mahajan S, Junttila M, et al. A transcriptional MAPK Pathway Activity Score (MPAS) is a clinically relevant biomarker in multiple cancer types. NPJ precision oncology. 2018;2(1):7. Epub 2018/06/07. doi: 10.1038/s41698-018-0051-4. PubMed PMID: 29872725; PubMed Central PMCID: PMCPMC5871852.

44. Foster R, Buckanovich RJ, Rueda BR. Ovarian cancer stem cells: Working towards the root of stemness. Cancer Lett. 2013;338(1):147–57. Epub 2012/11/10. doi: 10.1016/j.canlet.2012.10.023 S0304-3835(12)00628-3 [pii]. PubMed PMID: 23138176.

45. Roy L, Cowden Dahl KD. Can Stemness and Chemoresistance Be Therapeutically Targeted via Signaling Pathways in Ovarian Cancer? Cancers. 2018;10(8). Epub 2018/07/26. doi: 10.3390/cancers10080241. PubMed PMID: 30042330; PubMed Central PMCID: PMCPMC6116003.

46. Dhillon AS, Hagan S, Rath O, Kolch W. MAP kinase signalling pathways in cancer. Oncogene. 2007;26(22):3279–90. Epub 2007/05/15. doi: 10.1038/sj.onc.1210421. PubMed PMID: 17496922.

47. Liu S, Zha J, Lei M. Inhibiting ERK/Mnk/eIF4E broadly sensitizes ovarian cancer response to chemotherapy. Clinical & translational oncology: official publication of the Federation of Spanish Oncology Societies and of the National Cancer Institute of Mexico. 2018;20(3):374–81. Epub 2017/08/03. doi: 10.1007/s12094-017-1724-0. PubMed PMID: 28766096.

48. Gilmartin AG, Bleam MR, Groy A, Moss KG, Minthorn EA, Kulkarni SG, et al. GSK1120212 (JTP-74057) is an inhibitor of MEK activity and activation with favorable pharmacokinetic properties for sustained in vivo pathway inhibition. Clinical cancer research: an official journal of the American Association for Cancer Research. 2011;17(5):989–1000. Epub 2011/01/20. doi: 10.1158/1078-0432.ccr-10-2200. PubMed PMID: 21245089.

49. Yeh TC, Marsh V, Bernat BA, Ballard J, Colwell H, Evans RJ, et al. Biological characterization of ARRY-142886 (AZD6244), a potent, highly selective mitogen-activated protein kinase kinase 1/2 inhibitor. Clinical cancer research: an official journal of the American Association for Cancer Research. 2007;13(5):1576–83. Epub 2007/03/03. doi: 10.1158/1078-0432.ccr-06-1150. PubMed PMID: 17332304.

50. Hoeflich KP, Merchant M, Orr C, Chan J, Den Otter D, Berry L, et al. Intermittent administration of MEK inhibitor GDC-0973 plus PI3K inhibitor GDC-0941 triggers robust apoptosis and tumor growth inhibition. Cancer research. 2012;72(1):210–9. Epub 2011/11/16. doi: 10.1158/0008-5472.can-11-1515. PubMed PMID: 22084396.

51. Simpkins F, Jang K, Yoon H, Hew KE, Kim M, Azzam DJ, et al. Dual Src and MEK Inhibition Decreases Ovarian Cancer Growth and Targets Tumor Initiating Stem-Like Cells. Clinical cancer research: an official journal of the American Association for Cancer Research. 2018;24(19):4874–86. Epub 2018/07/01. doi: 10.1158/1078-0432.ccr-17-3697. PubMed PMID: 29959144.

52. Taniguchi T, Tischkowitz M, Ameziane N, Hodgson SV, Mathew CG, Joenje H, et al. Disruption of the Fanconi anemia-BRCA pathway in cisplatin-sensitive ovarian tumors. Nature medicine. 2003;9(5):568–74. Epub 2003/04/15. doi: 10.1038/nm852. PubMed PMID: 12692539.

53. Chang KE, Wei BR, Madigan JP, Hall MD, Simpson RM, Zhuang Z, et al. The protein phosphatase 2A inhibitor LB100 sensitizes ovarian carcinoma cells to cisplatin-mediated cytotoxicity. Molecular cancer therapeutics. 2015;14(1):90–100. Epub 2014/11/08. doi: 10.1158/1535-7163.mct-14-0496. PubMed PMID: 25376608; PubMed Central PMCID: PMCPMC4557740.

54. Haley J, Tomar S, Pulliam N, Xiong S, Perkins SM, Karpf AR, et al. Functional characterization of a panel of high-grade serous ovarian cancer cell lines as representative experimental models of the disease. Oncotarget. 2016;7(22):32810–20. Epub 2016/05/06. doi: 10.18632/oncotarget.9053. PubMed PMID: 27147568; PubMed Central PMCID: PMCPMC5078053.

55. Baba T, Convery PA, Matsumura N, Whitaker RS, Kondoh E, Perry T, et al. Epigenetic regulation of CD133 and tumorigenicity of CD133+ ovarian cancer cells. Oncogene. 2009;28(2):209–18. PubMed PMID: 18836486.

56. Curley MD, Therrien VA, Cummings CL, Sergent PA, Koulouris CR, Friel AM, et al. CD133 expression defines a tumor initiating cell population in primary human ovarian cancer. Stem cells. 2009;27(12):2875–83. Epub 2009/10/10. doi: 10.1002/stem.236. PubMed PMID: 19816957.

57. Kryczek I, Liu S, Roh M, Vatan L, Szeliga W, Wei S, et al. Expression of aldehyde dehydrogenase and CD133 defines ovarian cancer stem cells. Int J Cancer. 2012;130(1):29–39. Epub 2011/04/12. doi: 10.1002/ijc.25967. PubMed PMID: 21480217; PubMed Central PMCID: PMC3164893.

58. Choi YJ, Ingram PN, Yang K, Coffman L, Iyengar M, Bai S, et al. Identifying an ovarian cancer cell hierarchy regulated by bone morphogenetic protein 2. Proceedings of the National Academy of Sciences of the United States of America. 2015;112(50):E6882–8. Epub 2015/12/02. doi: 10.1073/pnas.1507899112. PubMed PMID: 26621735; PubMed Central PMCID: PMCPMC4687560.

59. Fremin C, Meloche S. From basic research to clinical development of MEK1/2 inhibitors for cancer therapy. Journal of hematology & oncology. 2010;3:8. Epub 2010/02/13. doi: 10.1186/1756-8722-3-8. PubMed PMID: 20149254; PubMed Central PMCID: PMCPMC2830959.

60. Meloche S, Pouyssegur J. The ERK1/2 mitogen-activated protein kinase pathway as a master regulator of the G1-to S-phase transition. Oncogene. 2007;26(22):3227–39. Epub 2007/05/15. doi: 10.1038/sj.onc.1210414. PubMed PMID: 17496918.

61. Zhao Y, Adjei AA. The clinical development of MEK inhibitors. Nature reviews Clinical oncology. 2014;11(7):385–400. Epub 2014/05/21. doi: 10.1038/nrclinonc.2014.83. PubMed PMID: 24840079.

62. Thigpen JT, Blessing JA, Ball H, Hummel SJ, Barrett RJ. Phase II trial of paclitaxel in patients with progressive ovarian carcinoma after platinum-based chemotherapy: a Gynecologic Oncology Group study. Journal of clinical oncology: official journal of the American Society of Clinical Oncology. 1994;12(9):1748–53. Epub 1994/09/01. doi: 10.1200/jco.1994.12.9.1748. PubMed PMID: 7916038.

63. Jordan MA, Toso RJ, Thrower D, Wilson L. Mechanism of mitotic block and inhibition of cell proliferation by taxol at low concentrations. Proceedings of the National Academy of Sciences of the United States of America. 1993;90(20):9552–6. Epub 1993/10/15. doi: 10.1073/pnas.90.20.9552. PubMed PMID: 8105478; PubMed Central PMCID: PMCPMC47607.

64. Takada M, Hix JML, Corner S, Schall PZ, Kiupel M, Yuzbasiyan-Gurkan V. Targeting MEK in a Translational Model of Histiocytic Sarcoma. Molecular cancer therapeutics. 2018;17(11):2439–50. Epub 2018/08/24. doi: 10.1158/1535-7163.MCT-17-1273. PubMed PMID: 30135215.

65. Sumi T, Hirai S, Yamaguchi M, Tanaka Y, Tada M, Niki T, et al. Trametinib downregulates survivin expression in RB1-positive KRAS-mutant lung adenocarcinoma cells. Biochemical and biophysical research communications. 2018;501(1):253–8. Epub 2018/05/05. doi: 10.1016/j.bbrc.2018.04.230. PubMed PMID: 29727601.

66. Vogel CJ, Smit MA, Maddalo G, Possik PA, Sparidans RW, van der Burg SH, et al. Cooperative induction of apoptosis in NRAS mutant melanoma by inhibition of MEK and ROCK. Pigment cell & melanoma research. 2015;28(3):307–17. Epub 2015/03/03. doi: 10.1111/pcmr.12364. PubMed PMID: 25728708.

67. Fernandez ML, DiMattia GE, Dawson A, Bamford S, Anderson S, Hennessy BT, et al. Differences in MEK inhibitor efficacy in molecularly characterized low-grade serous ovarian cancer cell lines. American journal of cancer research. 2016;6(10):2235–51. Epub 2016/11/09. PubMed PMID: 27822414; PubMed Central PMCID: PMCPMC5088288.

68. Sarvi S, Crispin R, Lu Y, Zeng L, Hurley TD, Houston DR, et al. ALDH1 Bio-activates Nifuroxazide to Eradicate ALDH(High) Melanoma-Initiating Cells. Cell chemical biology. 2018;25(12):1456–69.e6. Epub 2018/10/09. doi: 10.1016/j.chembiol.2018.09.005. PubMed PMID: 30293938; PubMed Central PMCID: PMCPMC6309505.

69. Zhang J, Guo X, Chang DY, Rosen DG, Mercado-Uribe I, Liu J. CD133 expression associated with poor prognosis in ovarian cancer. Modern pathology: an official journal of the United States and Canadian Academy of Pathology, Inc. 2012;25(3):456–64. Epub 2011/11/15. doi: 10.1038/modpathol.2011.170. PubMed PMID: 22080056; PubMed Central PMCID: PMCPMC3855345.

70. Liang J, Yang B, Cao Q, Wu X. Association of Vasculogenic Mimicry Formation and CD133 Expression with Poor Prognosis in Ovarian Cancer. Gynecologic and obstetric investigation. 2016;81(6):529–36. Epub 2016/05/11. doi: 10.1159/000445747. PubMed PMID: 27160772.

71. Stemberger-Papic S, Vrdoljak-Mozetic D, Ostojic DV, Rubesa-Mihaljevic R, Krigtofic I, Brncic-Fisher A, et al. Expression of CD133 and CD117 in 64 Serous Ovarian Cancer Cases. Collegium antropologicum. 2015;39(3):745–53. Epub 2016/02/24. PubMed PMID: 26898076.

72. Steg AD, Bevis KS, Katre AA, Ziebarth A, Dobbin ZC, Alvarez RD, et al. Stem Cell Pathways Contribute to Clinical Chemoresistance in Ovarian Cancer. Clin Cancer Res. 2012;18(3):869–81. Epub 2011/12/07. doi: 1078-0432.CCR-11-2188 [pii]10.1158/1078-0432.CCR-11-2188. PubMed PMID: 22142828.

73. Xia Y, Wei X, Gong H, Ni Y. Aldehyde dehydrogenase serves as a biomarker for worse survival profiles in ovarian cancer patients: an updated meta-analysis. BMC women’s health. 2018;18(1):199. Epub 2018/12/14. doi: 10.1186/s12905-018-0686-x. PubMed PMID: 30522488; PubMed Central PMCID: PMCPMC6284301.

