## Supplementary figures and images for "The MEK1/2 pathway as a therapeutic target in high-grade serous ovarian carcinoma"

### supplementary figures

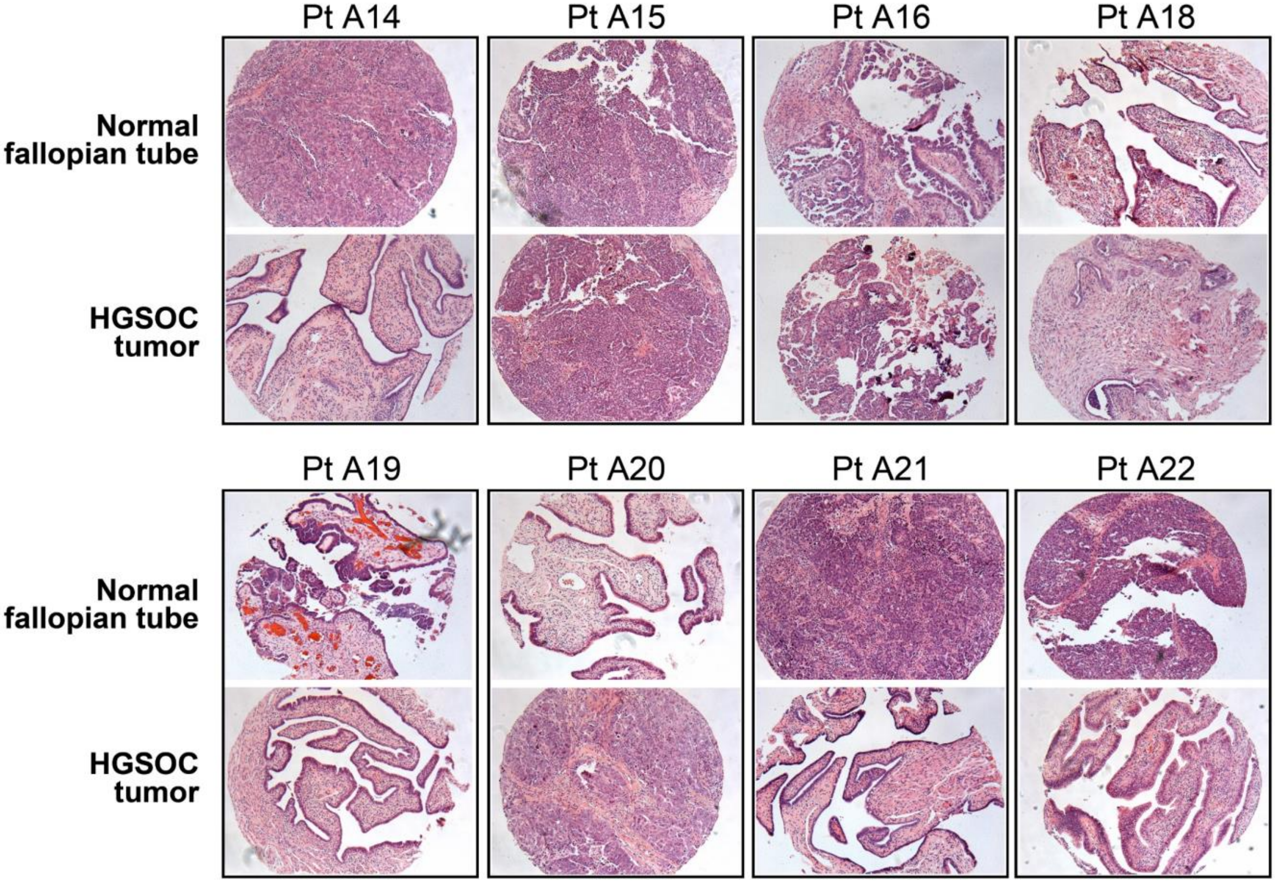

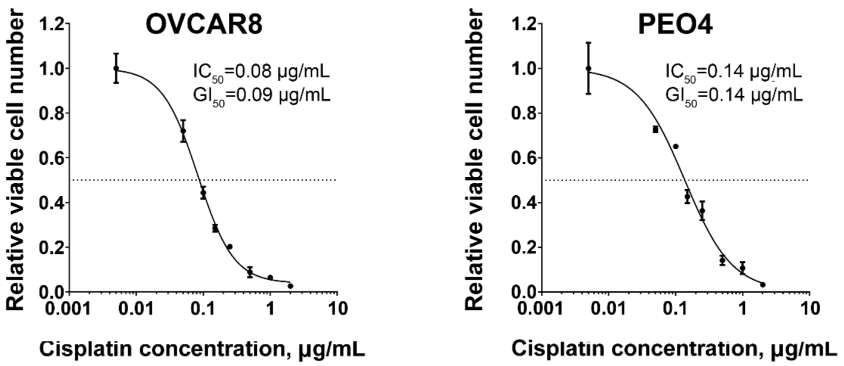

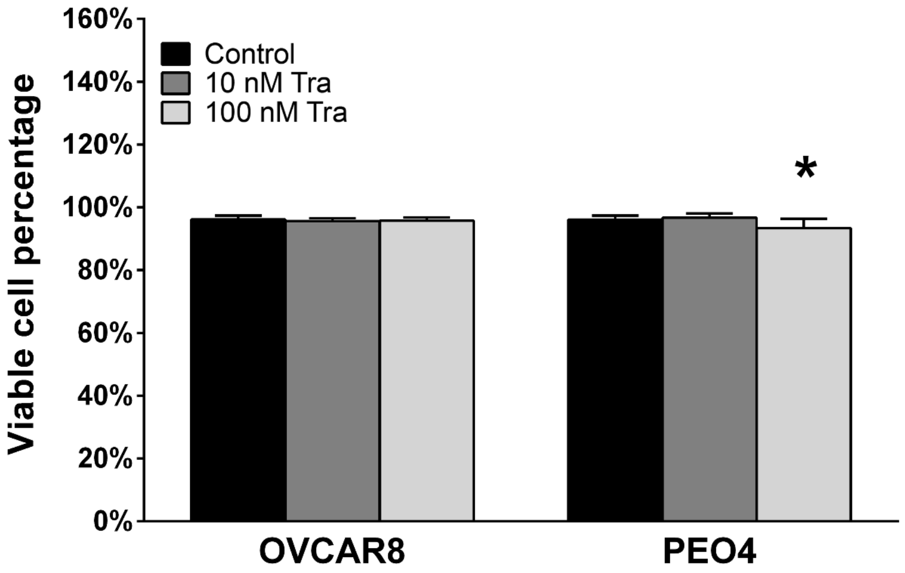

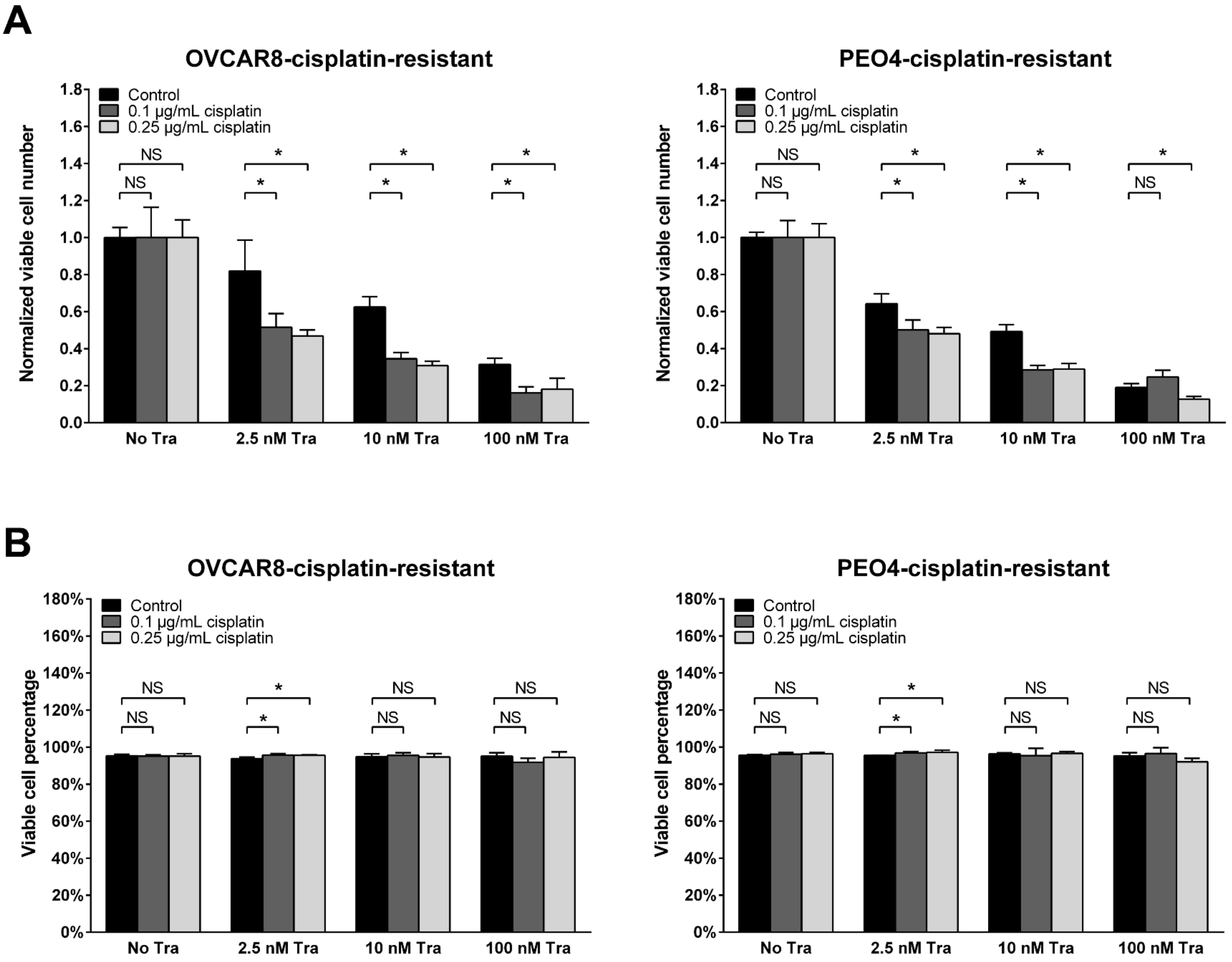

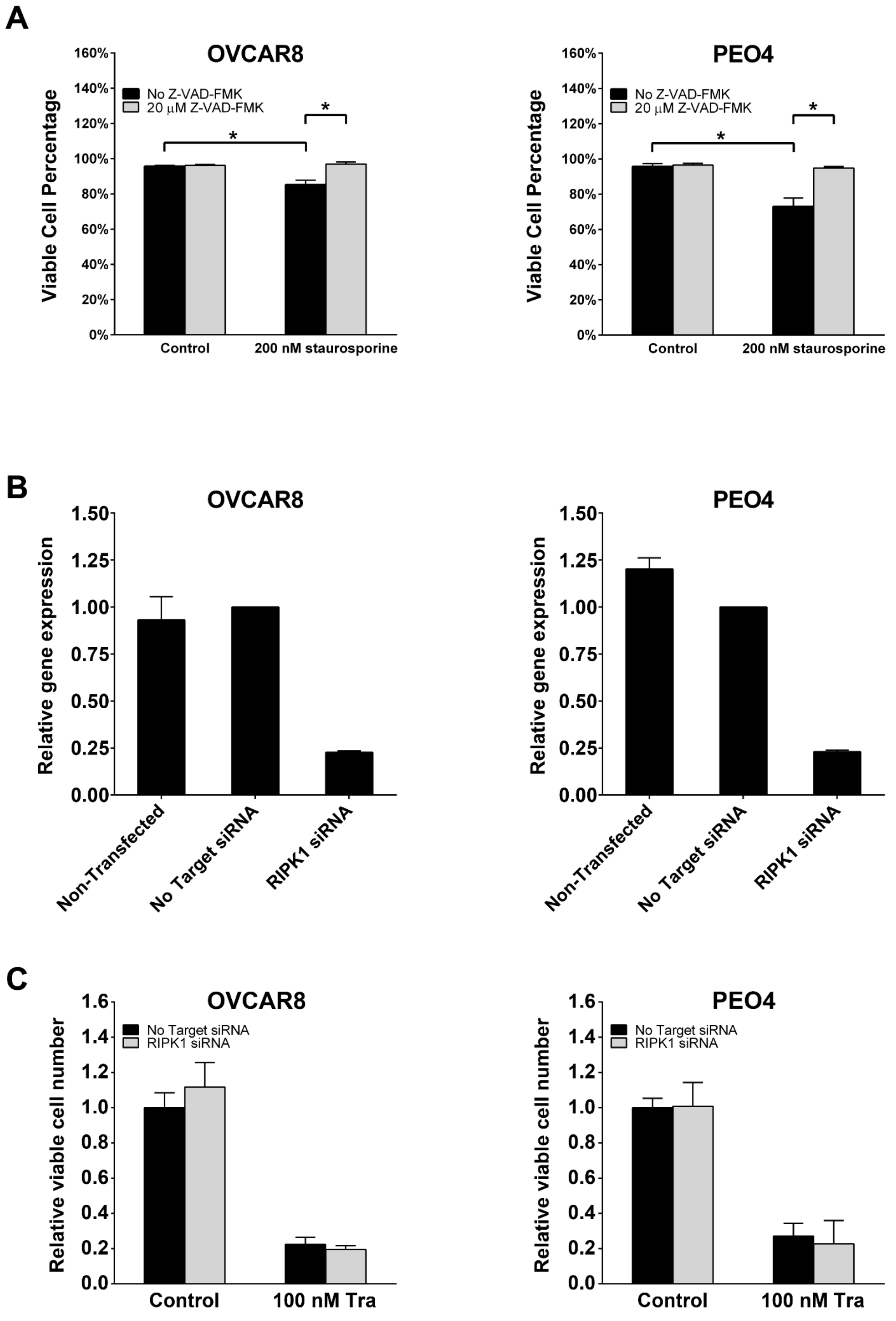

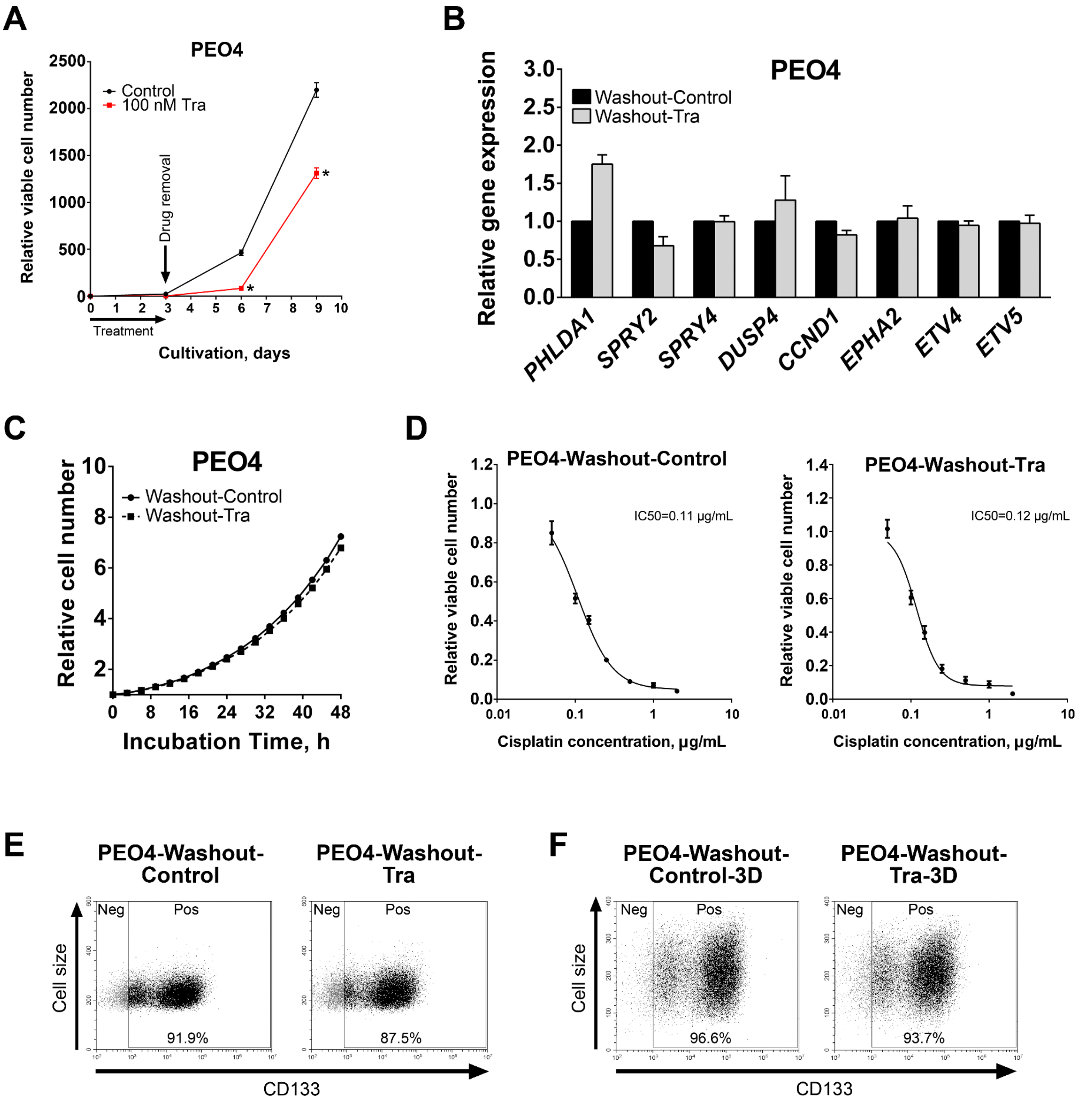
