## Supplementary material for "The MEK1/2 pathway as a therapeutic target in high-grade serous ovarian carcinoma": Suppl tables

**Table S1.** List of primers used in RT-PCR and RT-qPCR analysis.

| Gene | Forward primer | Reverse primer |
|---|---|---|
| TBP | GAGAGTTCTGGGATTGTACC | GGATTATATTCGGCGTTTCG |
| PHLDA1 | GGCAAGACAAGGTTTTGAGG | TCGCAAGTTTTCAGTAGGGTG |
| SPRY2 | TCCACTCAGCACAAACAC | GATTATGCCATCAGCAACAG |
| SPRY4 | CAACGGCTCTTAGACCAC | CACACTCCTTGCATTTACAC |
| DUSP4 | CCCCACTACACGACCAG | TCCGAGGAGACATTCAACAG |
| DUSP6 | GACGCTCGCTGTTTGTATC | GCTTCTAATCCCTCCCTCC |
| CCND1 | CTGGATGCTGGAGGTCTG | GGTCTCCTTCATCTTAGAGGC |
| EPHA2 | GTCAGCATCAACCAGACAGA | TCCCTTCTTGCGGTAAGTG |
| EPHA4 | CGGAGCGGAGAATGC | TCCTTCCCAGACAGAGTAG |
| ETV4 | AGGAGACATCAAGCAGGAAGGG | CCCAGAGCCTGGCGACC |
| ETV5 | AGCACAAGTTCCTGATGATG | CATAGTTAGCACCAAGAGCC |
| SOX2 | CACATGAACGGCTGGAG | CTGGTCATGGAGTTGTACTG |
| NANOG | AACTCTCCAACATCCTGAAC | GTAGGAAGAGTAAAGGCTGG |
| POU5F1 | GGGAAGGTATTCAGCCAAAC | AGAACCACACTCGGACC |
| KLF4 | CACACGGGATGATGCTC | GTCACAGTGGTAAGGTTTCT |
| ALDH1A1 | TGTTAGCTGATGCCGACTTG | TTCTTAGCCCGCTCAACACT |
| ALDH1A2 | CCATTGGAGTGTGTGGACAG | GATGAGGGCTCCCATGTAGA |
| ALDH1A3 | TCTCGACAAAGCCCTGAAGT | GTCCGATGTTTGAGGAAGGA |

**Table S2**. Main elements of MEK1/2-ERK1/2 signaling pathway.

| Protein name | UniProt ID | Gene name | NCBI Gene ID | Function | References |
|---|---|---|---|---|---|
| GRB2 | P62993 | *GRB2* | 2885 | Adaptor protein linking membrane receptors and RAS-controlled signaling pathways | [1-4] |
| SOS1 | Q07889 | *SOS1* | 6654 | Promoter of GDP-GTP exchange in RAS proteins | [5] |
| KRas<br>HRas<br>NRas | P01116<br>P01112<br>P01111 | *KRAS*<br>*HRAS*<br>*NRAS* | 3845<br>3265<br>4893 | Family of membrane-bound GTPases mediating signal transduction from GRB2 to RAF proteins | [6-9] |
| BRAF<br>RAF1 | P15056<br>P04049 | *BRAF*<br>*RAF1* | 673<br>5894 | Family of protein kinases activated by Ras proteins and targeting MEK1/2 | [7, 10-14] |
| MEK1<br>MEK2 | Q01986<br>P36507 | *MAP2K1*<br>*MAP2K2* | 5604<br>5605 | Protein kinases, central elements of MAP kinase signaling cascade targeting ERK1/2 | [10, 15, 16] |
| ERK1<br>ERK2 | P27361<br>P28482 | *MAPK3*<br>*MAPK1* | 5595<br>5594 | Protein kinases, main effectors of MAP kinase signaling cascade, phosphorylate more than 100 proteins | [10, 15, 17] |

**Table S3.** MEK-responsive gene signature described in [18].

| Gene name | NCBI Gene ID | Function | References |
|---|---|---|---|
| PHLDA1 | 22822 | Apoptosis regulator, may exert both pro- and anti-apoptotic effects in different tumors | [19, 20] |
| SPRY2 | 10253 | Proliferation suppressor, negative feedback regulator of MEK-ERK activity, promotes glioblastoma growth and chemoresistance | [21-23] |
| SPRY4 | 81848 | Proliferation suppressor, negative feedback regulator of MEK-ERK activity | [24, 25] |
| DUSP4 | 1846 | Proliferation and invasion suppressor, ERK phosphorylation inhibitor | [26, 27] |
| DUSP6 | 1848 | Proliferation suppressor, metastases formation suppressor, ERK phosphorylation inhibitor | [28, 29] |
| CCND1 | 595 | Proliferation stimulator, proto-oncogene | [30, 31] |
| EPHA2 | 1969 | Proliferation, EMT and invasion stimulator, apoptosis stimulator in some systems | [32-34] |

| | | | |
|---|---|---|---|
| EPHA4 | 2043 | Invasion stimulator, marker of poor prognosis | [35, 36] |
| ETV4 | 2118 | Proliferation and invasion stimulator, sustains cell stemness | [37-39] |
| ETV5 | 2119 | Proliferation stimulator, invasion suppressor, sustains cell stemness | [39-41] |


1. Belov AA, Mohammadi M. Grb2, a double-edged sword of receptor tyrosine kinase signaling. Science signaling. 2012;5(249):pe49. Epub 2012/11/08. doi: 10.1126/scisignal.2003576. PubMed PMID: 23131845; PubMed Central PMCID: PMCPMC3668340.
2. Braverman LE, Quilliam LA. Identification of Grb4/Nckbeta, a src homology 2 and 3 domain-containing adapter protein having similar binding and biological properties to Nck. The Journal of biological chemistry. 1999;274(9):5542-9. Epub 1999/02/20. PubMed PMID: 10026169.
3. Pandey P, Kharbanda S, Kufe D. Association of the DF3/MUC1 breast cancer antigen with Grb2 and the Sos/Ras exchange protein. Cancer research. 1995;55(18):4000-3. Epub 1995/09/15. PubMed PMID: 7664271.
4. Vindis C, Cerretti DP, Daniel TO, Huynh-Do U. EphB1 recruits c-Src and p52Shc to activate MAPK/ERK and promote chemotaxis. The Journal of cell biology. 2003;162(4):661-71. Epub 2003/08/20. doi: 10.1083/jcb.200302073. PubMed PMID: 12925710; PubMed Central PMCID: PMCPMC2173797.
5. Chardin P, Camonis JH, Gale NW, van Aelst L, Schlessinger J, Wigler MH, et al. Human Sos1: a guanine nucleotide exchange factor for Ras that binds to GRB2. Science (New York, NY). 1993;260(5112):1338-43. Epub 1993/05/28. PubMed PMID: 8493579.
6. Khan AQ, Kuttikrishnan S, Siveen KS, Prabhu KS, Shanmugakonar M, Al-Naemi HA, et al. RAS-mediated oncogenic signaling pathways in human malignancies. Seminars in cancer biology. 2018. Epub 2018/03/11. doi: 10.1016/j.semcancer.2018.03.001. PubMed PMID: 29524560.
7. Marshall M. Interactions between Ras and Raf: key regulatory proteins in cellular transformation. Molecular reproduction and development. 1995;42(4):493-9. Epub 1995/12/01. doi: 10.1002/mrd.1080420418. PubMed PMID: 8607981.
8. McCormick F. KRAS as a Therapeutic Target. Clinical cancer research : an official journal of the American Association for Cancer Research. 2015;21(8):1797-801. Epub 2015/04/17. doi: 10.1158/1078-0432.ccr-14-2662. PubMed PMID: 25878360; PubMed Central PMCID: PMCPMC4407814.
9. Nussinov R, Tsai CJ, Jang H. Oncogenic Ras Isoforms Signaling Specificity at the Membrane. Cancer research. 2018;78(3):593-602. Epub 2017/12/24. doi: 10.1158/0008-5472.can-17-2727. PubMed PMID: 29273632; PubMed Central PMCID: PMCPMC5811325.
10. Cseh B, Doma E, Baccarini M. "RAF" neighborhood: protein-protein interaction in the Raf/Mek/Erk pathway. FEBS letters. 2014;588(15):2398-406. Epub 2014/06/18. doi: 10.1016/j.febslet.2014.06.025. PubMed PMID: 24937142; PubMed Central PMCID: PMCPMC4099524.
11. von Kriegsheim A, Pitt A, Grindlay GJ, Kolch W, Dhillon AS. Regulation of the Raf-MEK-ERK pathway by protein phosphatase 5. Nature cell biology. 2006;8(9):1011-6. Epub 2006/08/08. doi: 10.1038/ncb1465. PubMed PMID: 16892053.
12. Roskoski R, Jr. RAF protein-serine/threonine kinases: structure and regulation. Biochemical and biophysical research communications. 2010;399(3):313-7. Epub 2010/08/03. doi: 10.1016/j.bbrc.2010.07.092. PubMed PMID: 20674547.

13. Brennan DF, Dar AC, Hertz NT, Chao WC, Burlingame AL, Shokat KM, et al. A Raf-induced allosteric transition of KSR stimulates phosphorylation of MEK. Nature. 2011;472(7343):366-9. Epub 2011/03/29. doi: 10.1038/nature09860. PubMed PMID: 21441910.
14. Migliaccio N, Sanges C, Ruggiero I, Martucci NM, Rippa E, Arcari P, et al. Raf kinases in signal transduction and interaction with translation machinery. Biomolecular concepts. 2013;4(4):391-9. Epub 2013/08/01. doi: 10.1515/bmc-2013-0003. PubMed PMID: 25436588.
15. Shaul YD, Seger R. The MEK/ERK cascade: from signaling specificity to diverse functions. Biochimica et biophysica acta. 2007;1773(8):1213-26. Epub 2006/11/23. doi: 10.1016/j.bbamcr.2006.10.005. PubMed PMID: 17112607.
16. Catling AD, Schaeffer HJ, Reuter CW, Reddy GR, Weber MJ. A proline-rich sequence unique to MEK1 and MEK2 is required for raf binding and regulates MEK function. Molecular and cellular biology. 1995;15(10):5214-25. Epub 1995/10/01. PubMed PMID: 7565670; PubMed Central PMCID: PMCPMC230769.
17. Yao Z, Seger R. The ERK signaling cascade--views from different subcellular compartments. BioFactors (Oxford, England). 2009;35(5):407-16. Epub 2009/07/01. doi: 10.1002/biof.52. PubMed PMID: 19565474.
18. Wagle MC, Kirouac D, Klijn C, Liu B, Mahajan S, Junttila M, et al. A transcriptional MAPK Pathway Activity Score (MPAS) is a clinically relevant biomarker in multiple cancer types. NPJ precision oncology. 2018;2(1):7. Epub 2018/06/07. doi: 10.1038/s41698-018-0051-4. PubMed PMID: 29872725; PubMed Central PMCID: PMCPMC5871852.
19. Murata T, Sato T, Kamoda T, Moriyama H, Kumazawa Y, Hanada N. Differential susceptibility to hydrogen sulfide-induced apoptosis between PHLDA1-overexpressing oral cancer cell lines and oral keratinocytes: role of PHLDA1 as an apoptosis suppressor. Experimental cell research. 2014;320(2):247-57. Epub 2013/11/26. doi: 10.1016/j.yexcr.2013.10.023. PubMed PMID: 24270013.
20. Neef R, Kuske MA, Prols E, Johnson JP. Identification of the human PHLDA1/TDAG51 gene: down-regulation in metastatic melanoma contributes to apoptosis resistance and growth deregulation. Cancer research. 2002;62(20):5920-9. Epub 2002/10/18. PubMed PMID: 12384558.
21. Yao Y, Luo J, Bian Y, Sun Y, Shi M, Xia D, et al. Sprouty2 regulates proliferation and survival of multiple myeloma by inhibiting activation of the ERK1/2 pathway in vitro and in vivo. Experimental hematology. 2016;44(6):474-82.e2. Epub 2016/03/27. doi: 10.1016/j.exphem.2016.02.009. PubMed PMID: 27016275.
22. Wang C, Delogu S, Ho C, Lee SA, Gui B, Jiang L, et al. Inactivation of Spry2 accelerates AKT-driven hepatocarcinogenesis via activation of MAPK and PKM2 pathways. Journal of hepatology. 2012;57(3):577-83. Epub 2012/05/24. doi: 10.1016/j.jhep.2012.04.026. PubMed PMID: 22617155; PubMed Central PMCID: PMCPMC3423481.
23. Walsh AM, Kapoor GS, Buonato JM, Mathew LK, Bi Y, Davuluri RV, et al. Sprouty2 Drives Drug Resistance and Proliferation in Glioblastoma. Molecular cancer research : MCR. 2015;13(8):1227-37. Epub 2015/05/03. doi: 10.1158/1541-7786.mcr-14-0183-t. PubMed PMID: 25934697; PubMed Central PMCID: PMCPMC4679183.
24. Zhou X, Xie S, Yuan C, Jiang L, Huang X, Li L, et al. Lower Expression of SPRY4 Predicts a Poor Prognosis and Regulates Cell Proliferation in Colorectal Cancer. Cellular physiology and biochemistry : international journal of experimental cellular physiology, biochemistry, and pharmacology. 2016;40(6):1433-42. Epub 2016/12/21. doi: 10.1159/000453195. PubMed PMID: 27997895.
25. Li M, Zhang H, Zhao X, Yan L, Wang C, Li C, et al. SPRY4-mediated ERK1/2 signaling inhibition abolishes 17beta-estradiol-induced cell growth in endometrial adenocarcinoma cell. Gynecological endocrinology : the official journal of the International Society of Gynecological Endocrinology. 2014;30(8):600-4. Epub 2014/05/09. doi: 10.3109/09513590.2014.912264. PubMed PMID: 24811094.
26. Ichimanda M, Hijiya N, Tsukamoto Y, Uchida T, Nakada C, Akagi T, et al. Downregulation of dual-specificity phosphatase 4 enhances cell proliferation and invasiveness in colorectal carcinomas. Cancer science. 2018;109(1):250-8. Epub 2017/11/19. doi: 10.1111/cas.13444. PubMed PMID: 29150975; PubMed Central PMCID: PMCPMC5765293.
27. Mazumdar A, Poage GM, Shepherd J, Tsimelzon A, Hartman ZC, Den Hollander P, et al. Analysis of phosphatases in ER-negative breast cancers identifies DUSP4 as a critical regulator of growth and invasion. Breast cancer research and treatment. 2016;158(3):441-54. Epub 2016/07/10. doi: 10.1007/s10549-016-3892-y. PubMed PMID: 27393618; PubMed Central PMCID: PMCPMC4963453.

28. Zhai X, Han Q, Shan Z, Qu X, Guo L, Zhou Y. Dual specificity phosphatase 6 suppresses the growth and metastasis of prostate cancer cells. Molecular medicine reports. 2014;10(6):3052-8. Epub 2014/09/23. doi: 10.3892/mmr.2014.2575. PubMed PMID: 25241655.
29. Okudela K, Yazawa T, Woo T, Sakaeda M, Ishii J, Mitsui H, et al. Down-regulation of DUSP6 expression in lung cancer: its mechanism and potential role in carcinogenesis. The American journal of pathology. 2009;175(2):867-81. Epub 2009/07/18. doi: 10.2353/ajpath.2009.080489. PubMed PMID: 19608870; PubMed Central PMCID: PMCPMC2716981.
30. Shan J, Zhao W, Gu W. Suppression of cancer cell growth by promoting cyclin D1 degradation. Molecular cell. 2009;36(3):469-76. Epub 2009/11/18. doi: 10.1016/j.molcel.2009.10.018. PubMed PMID: 19917254; PubMed Central PMCID: PMCPMC2856324.
31. Casimiro MC, Velasco-Velazquez M, Aguirre-Alvarado C, Pestell RG. Overview of cyclins D1 function in cancer and the CDK inhibitor landscape: past and present. Expert opinion on investigational drugs. 2014;23(3):295-304. Epub 2014/01/07. doi: 10.1517/13543784.2014.867017. PubMed PMID: 24387133.
32. Wen Q, Chen Z, Chen Z, Chen J, Wang R, Huang C, et al. EphA2 affects the sensitivity of oxaliplatin by inducing EMT in oxaliplatin-resistant gastric cancer cells. Oncotarget. 2017;8(29):47998-8011. Epub 2017/06/19. doi: 10.18632/oncotarget.18208. PubMed PMID: 28624791; PubMed Central PMCID: PMCPMC5564621.
33. Chen P, Huang Y, Zhang B, Wang Q, Bai P. EphA2 enhances the proliferation and invasion ability of LNCaP prostate cancer cells. Oncology letters. 2014;8(1):41-6. Epub 2014/06/25. doi: 10.3892/ol.2014.2093. PubMed PMID: 24959216; PubMed Central PMCID: PMCPMC4063646.
34. Zhang G, Njauw CN, Park JM, Naruse C, Asano M, Tsao H. EphA2 is an essential mediator of UV radiation-induced apoptosis. Cancer research. 2008;68(6):1691-6. Epub 2008/03/15. doi: 10.1158/0008-5472.can-07-2372. PubMed PMID: 18339848; PubMed Central PMCID: PMCPMC4469360.
35. Miyazaki K, Inokuchi M, Takagi Y, Kato K, Kojima K, Sugihara K. EphA4 is a prognostic factor in gastric cancer. BMC clinical pathology. 2013;13(1):19. Epub 2013/06/07. doi: 10.1186/1472-6890-13-19. PubMed PMID: 23738943; PubMed Central PMCID: PMCPMC3720259.
36. Liu C, Huang H, Wang C, Kong Y, Zhang H. Involvement of ephrin receptor A4 in pancreatic cancer cell motility and invasion. Oncology letters. 2014;7(6):2165-9. Epub 2014/06/17. doi: 10.3892/ol.2014.2011. PubMed PMID: 24932309; PubMed Central PMCID: PMCPMC4049679.
37. Qi M, Liu Z, Shen C, Wang L, Zeng J, Wang C, et al. Overexpression of ETV4 is associated with poor prognosis in prostate cancer: involvement of uPA/uPAR and MMPs. Tumour biology : the journal of the International Society for Oncodevelopmental Biology and Medicine. 2015;36(5):3565-72. Epub 2014/12/30. doi: 10.1007/s13277-014-2993-7. PubMed PMID: 25544710.
38. Moss AC, Lawlor G, Murray D, Tighe D, Madden SF, Mulligan AM, et al. ETV4 and Myeov knockdown impairs colon cancer cell line proliferation and invasion. Biochemical and biophysical research communications. 2006;345(1):216-21. Epub 2006/05/09. doi: 10.1016/j.bbrc.2006.04.094. PubMed PMID: 16678123.
39. Akagi T, Kuure S, Uranishi K, Koide H, Costantini F, Yokota T. ETS-related transcription factors ETV4 and ETV5 are involved in proliferation and induction of differentiation-associated genes in embryonic stem (ES) cells. The Journal of biological chemistry. 2015;290(37):22460-73. Epub 2015/08/01. doi: 10.1074/jbc.M115.675595. PubMed PMID: 26224636; PubMed Central PMCID: PMCPMC4566222.
40. Llaurado M, Abal M, Castellvi J, Cabrera S, Gil-Moreno A, Perez-Benavente A, et al. ETV5 transcription factor is overexpressed in ovarian cancer and regulates cell adhesion in ovarian cancer cells. International journal of cancer. 2012;130(7):1532-43. Epub 2011/04/27. doi: 10.1002/ijc.26148. PubMed PMID: 21520040.
41. Monge M, Colas E, Doll A, Gil-Moreno A, Castellvi J, Diaz B, et al. Proteomic approach to ETV5 during endometrial carcinoma invasion reveals a link to oxidative stress. Carcinogenesis. 2009;30(8):1288-97. Epub 2009/05/16. doi: 10.1093/carcin/bgp119. PubMed PMID: 19443906.
